# SpyDen: Automating molecular and structural analysis across spines and dendrites

**DOI:** 10.1101/2024.06.07.597872

**Authors:** Maximilian F. Eggl, Surbhit Wagle, Jean P. Filling, Thomas E. Chater, Yukiko Goda, Tatjana Tchumatchenko

**Affiliations:** Institute of Neuroscience CSIC-UMH, Alicante, Spain; Institute of Experimental Epileptology and Cognition Research, University of Bonn Medical centre, Venusberg-Campus 1, 53127 Bonn, Germany; University of Mainz Medical centre, Anselm-Franz-von-Bentzel-Weg 3, 55128 Mainz, Germany; Laboratory for Synaptic Plasticity and Connectivity, RIKEN centre for Brain Science, Wako-shi, Saitama, Japan; Department of Physiology, Keio University School of Medicine, Tokyo, Japan; Synapse Biology Unit, Okinawa Institute of Science and Technology Graduate University, Onna-son, Kunigami-gun, Okinawa, Japan

## Abstract

Investigating the molecular composition across neural compartments such as axons, dendrites, or synapses is critical for our understanding of learning and memory. State-of-the-art microscopy techniques can now resolve individual molecules and pinpoint their position with micrometre or even nanometre resolution across tens or hundreds of micrometres, allowing the labelling of multiple structures of interest simultaneously. Algorithmically, tracking individual molecules across hundreds of micrometres and determining whether they are inside any cellular compartment of interest can be challenging. Historically, microscopy images are annotated manually, often using multiple software packages to detect fluorescence puncta (e.g. labelled mRNAs) and then trace and quantify cellular compartments of interest. Advanced ANN-based automated tools, while powerful, are often able to help only with selected parts of the data analysis pipeline, may be optimised for specific spatial resolutions or cell preparations or may not be fully open source and open access to be sufficiently customisable. To address these challenges, we developed SpyDen. SpyDen is a Python package based upon three principles: *i)* ease of use for multi-task scenarios, *ii)* open-source accessibility and data export to a common, open data format, *iii)* the ability to edit any software-generated annotation and generalise across spatial resolutions. Equipped with a graphical user interface and accompanied by video tutorials, SpyDen provides a collection of powerful algorithms that can be used for neurite and synapse detection as well as fluorescent puncta and intensity analysis. We validated SpyDen using expert annotation across numerous use cases to prove a powerful, integrated platform for efficient and reproducible molecular imaging analysis.

## Introduction

Neuronal synapses and other cell compartments contain molecules that are critical for the implementation of information processing in the brain (Abbott and Regehr, 2004). Moreover, synaptic plasticity, the ability of synapses, especially that of spines, to change in size and strength based on computational demands, is important for learning and overall brain function and is synthesised through a complex interaction of different molecular players (Yau, 1976; Rosahl et al., 1993; Zilberter, 2000; Yuste and Bonhoeffer, 2001; Chevaleyre et al., 2006; Abbott and Regehr, 2004; Lüscher and Malenka, 2012; Frémaux and Gerstner, 2016; Marblestone et al., 2016; Eggl et al., 2023; Wagle et al., 2023). Recent microscopy advances (Van Harreveld and Fifkova, 1975; Desmond and Levy, 1986; Hosokawa et al., 1995; Matsuzaki et al., 2004; Kwon and Sabatini, 2011; Nägerl and Bonhoeffer, 2010; Ding et al., 2009; Schermelleh et al., 2019; Padmanabhan et al., 2021) have enabled the study of the complex interactions between synaptic plasticity and neural function and the underlying molecular dynamics taking place over time across the dendritic tree, axons and synapses at micrometre to nanometre scale. Additionally, these novel techniques allow for the quantification of the localisation profiles of mRNAs and different protein species along the dendritic or axonal tree as well as within individual synapses, thereby providing insight into neuronal synaptic and dendritic dynamics at unprecedented resolution (Holt and Schuman, 2013; Rangaraju et al., 2017; Helm et al., 2021). One bottleneck to the analysis of these datasets is manual or semi-manual annotation (commonly with *Neuronstudio* (Rodriguez et al., 2008), java-based *Fiji* (Schindelin et al., 2012) (for example Gilles et al., 2024) or python based *NAPARI* Ahlers et al. (2023)), which can be a time-consuming and labour-intensive task. The analysis can also be confounded by the observation that even experienced human annotators show a significant amount of variation (Fernholz et al., 2023; Graves et al., 2021). Additionally, the variability of neural anatomy (Mishchenko et al., 2010; Mukai et al., 2011; Berry and Nedivi, 2017; Graves et al., 2021) and the challenges of tracking structures of interest across space and time (Fernholz et al., 2023) indicate the need for more quantitative and comprehensive analysis tools that can accommodate the diversity of cell compartments.

The increasing amount of high-resolution imaging data, coupled with the above-mentioned challenges, means that automatic approaches are becoming more pragmatic for data analysis (Das et al., 2021; Ghani et al., 2017). With the application of deep neural networks, the opportunity to perform automatic and reproducible analysis at a level similar to human-level annotation is within reach(Tong et al., 2021; Vogel et al., 2023; Ekaterina et al., 2023). A common approach is to train artificial neural networks (often U-nets) to segment spines and dendrites in 3D from 2D stacked images and to generate 3D meshes (Ronneberger et al., 2015; Vidaurre-Gallart et al., 2022). Other artificial neural net approaches include the Matlab package *SpineS* (Argunşah et al., 2022), the DeepD3 Framework (Fernholz et al., 2023), or the work of Vogel et al. (2023). All these approaches represent powerful tools for the analysis of dendritic stretches and offer the ability to utilise a pre-trained ANN or train a personal network.

However, several challenges can arise when applying these approaches. First, training deep learning models on custom datasets is time and energy-consuming and requires machine learning knowledge. Second, the resulting segmentations can require further processing to extract data of interest, e.g. centres and ROIs of spines, distance of spines from soma or the medial axis path of the dendrite or number of molecules of interest within a cell compartment (Iannella and Tanaka, 2006; Kastellakis et al., 2015; Chater et al., 2022). Third, the resulting ROIs (dendrite and spines) are often not editable once calculated by the underlying algorithms. Providing a way to interact with the automatically generated results is critical for effective and robust analysis if the underlying algorithm produces erroneous results in specific image areas. Fourth, some of the available tools require continuous, spatial signals, while others work with discrete puncta data and apply different definitions of cell compartments, making comparisons difficult. This makes a comprehensive and reproducible data analysis across different molecular data sets challenging.

To close this gap, we present SpyDen, an open-access and open-source package to automatically evaluate and trace multiple features of interest, including dendrites and spines, as well as mRNA and protein localisations. This work includes a brief overview of the SpyDen workflow, a description of the underlying algorithms and a set of validations performed on a variety of datasets. Given this, we believe that SpyDen opens up new avenues for image analysis which are both time-efficient and reproducible in the study of dendrites and spines as well as beyond.

### Data analysis with SpyDen

We designed SpyDen as an integrated platform with the following properties

- Reliable, cross-validated algorithms for the analysis of dendritic segments, somata and spines that provide both the ROIs and relevant statistics across space and time that are bench-marked across different image resolutions using human expert performance.
- The integrated analysis of continuous molecular signals as well as discrete puncta that represent individual molecules across structures of interest means that comparisons between cell compartments and data types are possible within a single user interface and sets of parameters.
- A graphical user interface (see Fig. S1) that can be used without any prior programming knowledge. By providing a compiled executable, scientists can utilise SpyDen without the need to read and write computer code. To further enhance the user experience, we provide video tutorials.
- SpyDen is fully open-source and open-access, containing no proprietary software. Its algorithms can be further customised to serve alternative analysis goals both within and outside the neuroscience context.
- All results from the automatic algorithms can be edited, both at the algorithm level and the local ROI levels. This means that SpyDen provides a robust, algorithm-provided baseline that can be customised further if necessary, a feature not available in many current software tools.

SpyDen combines a set of powerful algorithms within a single GUI that is optimised for ease of use across different analysis scenarios. This allows for efficient and reproducible analysis of neuronal images, avoiding the need to switch to different programs for different steps, reducing the number of potential errors, and speeding up image analysis. See Fig. 1 for an example of a complete workflow. Next, we detail the SpyDen workflow to demonstrate the above features and analysis steps, showing how SpyDen can be employed to tackle complex biological imaging challenges and ensure precision and ease of use.

**Figure 1:**
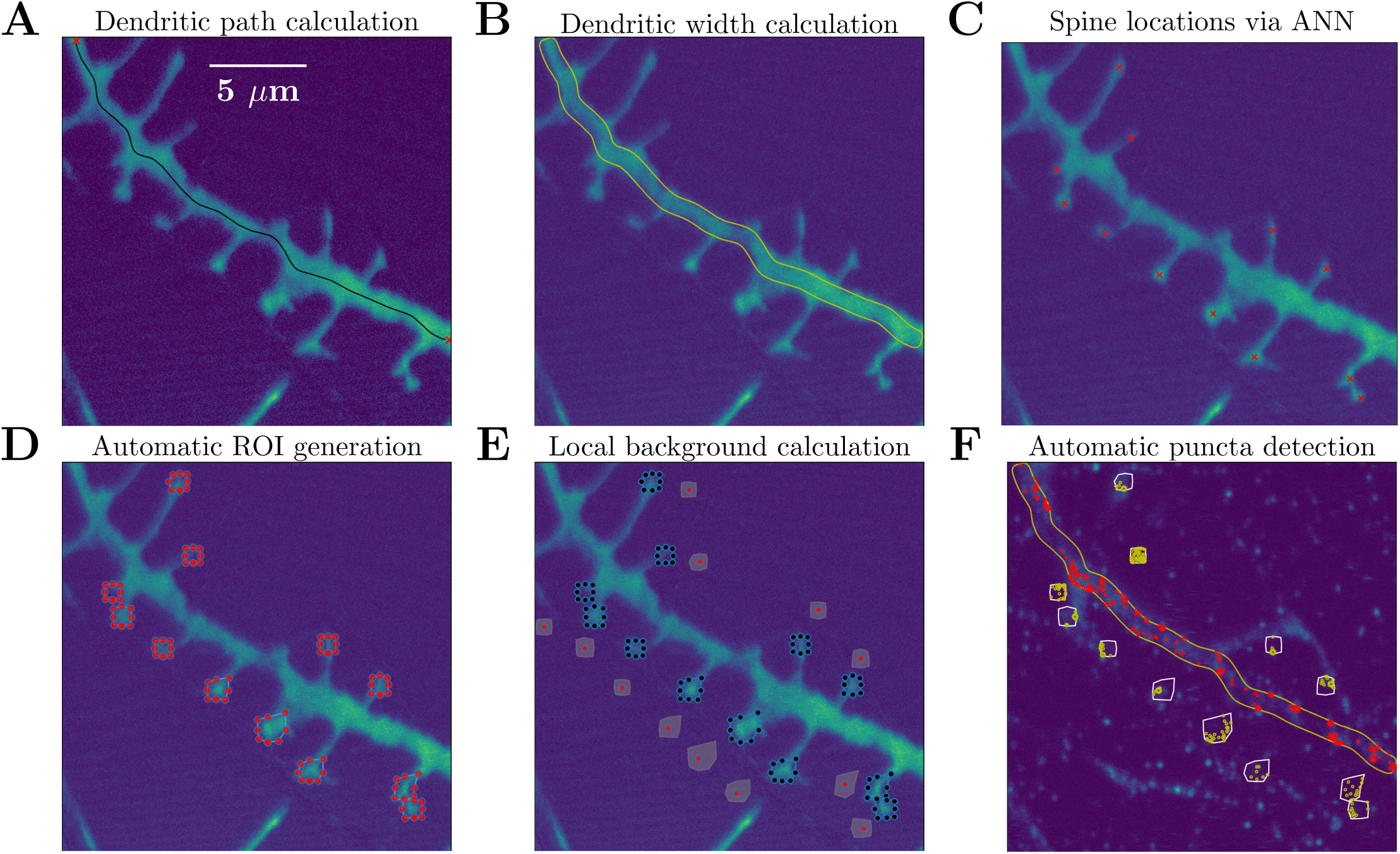
Steps of SpyDen’s data analysis pipeline illustrated with an example experimental image. **A)** Result of the medial axis calculation. The two red markers depict the user selection and the black line is the path through the dendrite connecting those points. **B)** The width of the dendrite, using the medial axis, is calculated and depicted in yellow. **C)** The red crosses mark the spine locations, obtained here through the use of a neural network (manual selection can also be performed). **D)** Using the spine locations, editable ROIs (depicted by the polygons with the red vertices) are calculated. **E)** To obtain statistics on the local background, the user can move the background location (depicted by the red markers surrounded by the grey). **F)** Using the dendritic ROI (red points, from *B*) and ROIs from the spines (yellow points, calculated in *D*) puncta can be calculated for fluorescence signal detection.

### Analysing the dendritic segments

The first component in the SpyDen analysis pipeline is the generation of medial axis paths using user-provided start and end points that best describe a set of chosen dendrites. The resulting medial axis path (or paths) are available not only for dendrite analysis but can also be used for other parts of the analysis pipeline, e.g. spine analysis. An example of a medial axis path is shown in Fig. 1A, where the horizontal dendrite path has been generated (black line) connecting the two marked end-points (red crosses).

In line with our goal that all results arising from SpyDen algorithms should be editable, we provide the users the possibility to modify this path if desired (i) allowing for the selection of the channel that depicts the dendrite most reliably using the associated SpyDen sliders and (ii) including slider settings determining the level of background noise filtering. An example of noise filtering can be seen in Fig. 2A or for more details see Supplementary Fig. S2. While the above strategy allows for the global enhancement of the path, we also offer the users the ability to edit local parts of the path. Thus, we have included interactable vertices along the medial axis path (see Fig. 2A). These fully editable nodes (nodes can be moved, deleted, or new ones added) allow for fine adjustments to be made to single points of the path.

**Figure 2:**
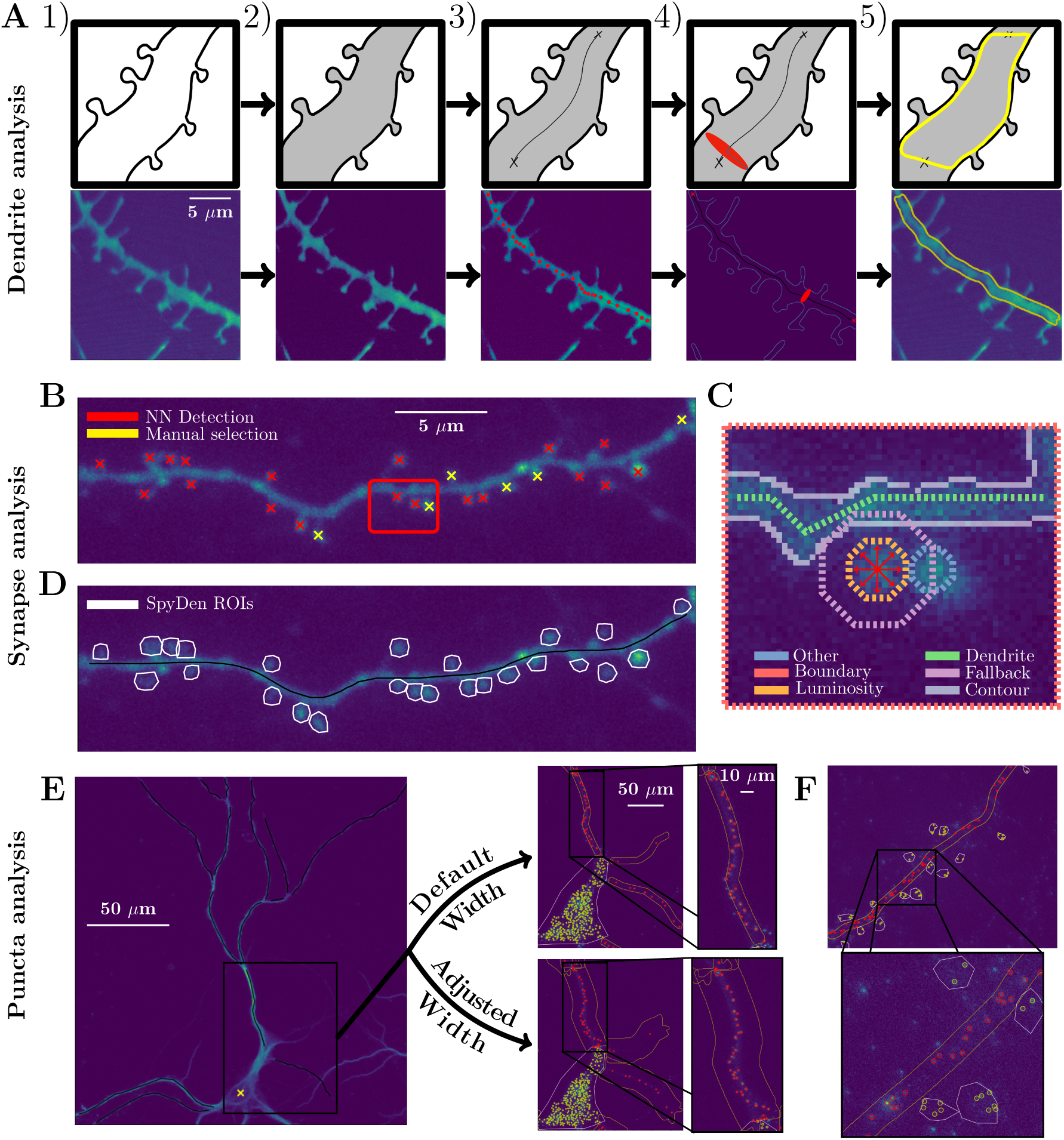
Illustrations of the three analysis pipelines of SpyDen with example images. **A)** Analysis pipeline for generating the dendritic segmentation. From left to right): 1) Input image of the dendrite and the dendritic spines. 2) A median filter is applied to reduce the salt and pepper noise followed by a chosen threshold (default is the mean) for a rough segmentation of the fore- and background. 3) Given the start and end point of the dendrite, the medial axis path will be calculated. 4) For every point on the medial axis path we fit an ellipse that covers the dendrite to determine its width. 5) Applying a width increase condition, we require continuity in the dendritic width calculation and do not allow large abrupt changes that can occur due to the presence of spines. **B)** An example dendritic stretch, where the spine heads have been marked manually (yellow) as well as using an artificial neural network (red). The red box is additionally shown in a zoomed-in version in C). **C)** Using the selected centre of the spine (centre of the red arrows), we project outwards in eight directions until a certain number of conditions are satisfied. The conditions are depicted with dashed lines in different colours on the plot. Example conditions are the dendrite rule (spine ROI should not intersect with the dendrite) or the fall-back rule (the spine ROI should exceed a certain threshold). **D)** After iterating through all spine centres, and applying the heuristic rules to generate spine ROIs (white lines), we are left with a set of potential ROIs that can be further altered if necessary. **E-F)** Once the ROIs (Soma, dendritic width - in *E* - and spine - in *F*) are finalised, puncta detection can be used to find bright spot puncta in the ROIs and get various measures for each punctum. Dendritic segmentation needs to be performed to obtain the dendritic puncta.

With the medial axis path calculation results, SpyDen can then generate the full dendritic segmentation (see yellow outline in Fig. 1B). Once again, the result of this automatic process can be enhanced by using a set of sliders that alter the behaviour of the underlying segmentation algorithm. Examples of the segmentation results after altering these sliders can be seen in Supplementary Fig. S2F-H.

At this point, the results of the dendritic analysis can be saved in *.json* and *.csv* files (which are both human and machine-readable). These files include statistics such as the width of the dendrite, luminosity along the medial axis and luminosity of the segmentation result (see table S1). The masks for dendritic segmentation for each dendrite are also saved as images. Additionally, the medial axis and dendrite segmentation ROIs are saved as *.png* and *.roi* files that either can be used for further post-processing or directly loaded into *ImageJ*, respectively.

### Detecting and analysing spines

Even though it has been demonstrated that even experienced human annotators show significant variation when detecting and analysing spines (Fernholz et al., 2023; Graves et al., 2021), many experimental labs still rely on manual analysis. To overcome this challenge and provide a baseline to reduce this variation, SpyDen combines automatic and manual approaches to provide spine statistics. On the one hand, a neural network-based implementation (see Supplementary Fig. S4 for a schematic) can be used for initial detection, which is then augmented by directly clicking on the spines (see Fig. 2B). Additional functionality is provided beyond merely selecting the spines: spine selections (both manual and automatic) can be deleted, and spines of particular interest (for example, spines previously targeted for stimulation) can also be highlighted. This highlighting provides additional flexibility with regard to other neuronal structures, such as neuronal somata, that can be selected for analysis as if they were spines.

Given the set of spine locations, SpyDen then sequentially generates ROIs surrounding those structures. The ROI generation is based on a set of heuristic rules (for a depiction of these rules applied to an example spine, see Fig. 2C) that take advantage of the inherent structure of the spines and rely on a set of parameters that can be tuned to achieve results that better fit the chosen experimental paradigm. We demonstrate the difference in ROI generation when different parameters are chosen by interacting with the sliders provided by SpyDen in Supplementary Fig. S5. Similarly to the medial axis path generated for the dendritic branch, the spine ROIs are made up of interactable vertices (see Fig. 2D), which means that the automatic results can also be edited on a spine-to-spine level, enhancing the results.

SpyDen additionally provides a way to calculate the local background luminosity of each spine. Given that the local luminosity of the image surrounding the spine may impact its relative brightness and thus affect the resulting analysis, providing these background values is critical for accurate results. Therefore, SpyDen calculates a location close to each spine and uses the associated ROI to calculate the image’s local background luminosity (see Fig. 1E). Within the design philosophy of SpyDen, the location of this background can be manually adjusted if the algorithm chooses an unsuitable location.

A further challenge when analysing spines in temporal data is that the images may shift both on a global (on an image level if, for example, the sample was disturbed) or on a local level (the microscopic movements of the spines or dendrites themselves). To combat this challenge, SpyDen provides both global and local motion correction and allows for accurate measurements. In the case of the local correction, manual intervention is once more provided by selecting a given timestep and dragging the associated ROI to the correct location.

Once all the above steps are completed, the spine statistics are stored as *.json* and *.csv* files and contain a wide variety of metrics for further post-processing. Additionally, *.png* and *.roi* files are generated of the spine polygons, which can be used for further analysis or figure generation.

### Detecting and analysing fluorescent puncta inside structures of interest

Finally, SpyDen takes advantage of its all-in-one nature to use the previous analysis (spatial image information) to detect and measure fluorescent puncta (discrete datapoints) inside the generated ROI.

As we have two types of ROIs (polygonal spine ROIs - which also can be used to study soma and other similar structures- and segmented line ROIs for dendrites), we note that puncta are detected differently for each of them. For polygonal ROIs, fluorescent puncta are detected inside the area enclosed by the polygon (yellow points, Fig. 1F). For segmented line ROIs, the fluorescent puncta are detected by considering local rectangles defined by the cross-section and the closest line segments (red points, Fig. 1F). SpyDen provides the capability to filter out puncta of certain minimum and maximum sizes and omits puncta whose intensity is below the background noise level. This improves the efficiency of the algorithm and avoids detecting spurious puncta. The coordinates of the detected puncta are saved in human and machine-readable *.json* and *.csv* files.

## SpyDen Methods

### Algorithms for dendrite analysis

#### Medial axis path calculation

To obtain the optimal medial axis that sits in the middle of the selected dendrite, we consider the treatment of the experimental image as an optimal path problem, i.e., given the start and end points provided, we attempt to find the best path through the dendrite. The image is transformed into a binary matrix, where the pixels that have luminosity over a given threshold represent an admissible path, and all other pixels are inaccessible (walls) (see Fig. S2B). Given this formulation, standard pathing algorithms can be applied to obtain the shortest path calculation. SpyDen utilises the standard Dijkstra algorithm of NetworkX (Hagberg et al., 2008) and speeds up the calculation time by downsampling the image if it is larger than 512×512 pixels.

However, if we solely applied the algorithm to the matrix described above, the path we generate would not lie within the middle of the dendrite but instead favour the dendrite edges (especially if the dendrite is curved or angled). Therefore, we augment the binary matrix, which represents the admissible dendrite pixels, with another factor that represents how far a given pixel is from the boundary of the dendrite. This factor is calculated by checking how far a given pixel is from an inadmissible pixel (i.e., a pixel below the luminosity threshold), i.e., a simple approximation of how far a given pixel is from the edge of the dendrite. This alteration ensures that the algorithm generates the shortest path at the dendrite’s centre. For an example of this augmented formulation, see Fig. S2C, where the peaks represent the points furthest from the dendrite edge.

The resulting path consists of all pixels that connect the dendritic endpoints. Retaining all of these pixels, however, is inefficient, makes saving the dendrite for future use difficult, and complicates the editability of the dendritic path. To overcome these problems, the full-rank path is compressed by removing redundant pixels and generating a path consisting of a smaller set of control points. These control points, which define the editable nodes seen in Figure 2A, are major turning points of the path and are obtained by a curvature-dependent sampling of the original path. By interpolating linearly between them, the full rank path can be recovered. A pseudo-implementation of the above-described algorithm can be found in the supplemental material in Algorithm 1.

#### Dendritic width calculation

Given the full-rank medial axis path, we can then proceed to segment the selected stretch of the dendrite in its entirety. To determine the exact width of the dendrite at each point, we define an ellipse at each pixel of the full-rank medial axis path. The semi-minor axis of these ellipses is set to a pre-set small value and points in the direction along the medial axis. In contrast, the semi-major axis is normal to the medial axis and is iteratively increased until it intersects with the boundary of the dendrite. An example illustration of an ellipse can be seen in the top panel of Figure 2A. This dendrite boundary is calculated using the Canny edge detection algorithm (Canny, 1986) applied to the median filtered and thresholded image. We employ this ellipse-based approach to generate the dendrite segmentation for two reasons: Firstly, directly using the canny edge detection detects *all* edges in the image, meaning that synaptic edges are not differentiated from the dendritic boundary. Secondly, depending on the luminosity profile of the image, certain edges may not be detected, leading to gaps in the dendritic boundary. By using ellipses and increasing their width iteratively, we are guaranteed to intersect with an edge at some point, which is not necessarily true if we solely used an outward-pointing ray. Examples of these phenomena can be seen in Fig. S3.

To ensure that the dendritic width calculation does not include structures growing out of the dendrite (i.e. filopodia, spines, etc.), the algorithm includes a smoothing condition that avoids abrupt changes in the width. To provide a means to interact with this algorithm, SpyDen includes two sliders that alter (i) the width-multiplication factor and (ii) the effect of the smoothing condition. These two sliders allow for significant alterations to the segmentation result, which in turn can enhance the result of the dendritic width calculation. The full algorithmic description of the dendritic width calculation can be seen in Algorithm 2.

### Synaptic analysis

#### Neural network-based identification of spine heads

Artificial neural networks are an effective tool to automatically annotate spines along dendritic stretches for further analysis (Ronneberger et al., 2015; Kashiwagi et al., 2019; Tong et al., 2021; Argunşah et al., 2022; Fernholz et al., 2023; Vogel et al., 2023). In most cases, this approach is employed directly for segmentation rather than the localisation of spines along the dendritic branches. When spines are segmented directly, any additional ones not identified in the segmentation process can be difficult to include in later steps of the data-analysis pipeline. Additionally, most approaches mentioned above cannot edit and correct the generated ROIs once they are calculated.

These limitations mean that we opted to use a faster R-CNN architecture (Girshick, 2015) to locate the centre of mass of spines directly (similar to the approach taken to Vogel et al., 2023) rather than segmenting them. Concretely, the fast R-CNN architecture takes in an image of variable size and outputs a list of spine-bounding boxes and associated confidences. Given the centre of these boxes, we can then calculate the spine’s centre. Sets of spines from three different datasets (Helm et al., 2021; Chater et al., 2022; Fernholz et al., 2023) were manually labelled and used to train the neural network (pre-trained on ImageNet). Additional training images were generated by altering the original training set by changing their resolutions and rotating or mirroring them. For a set of example training images, as well as the schematic of the architecture, see supplemental figure Fig. S4.

Focusing on locating spines, rather than segmenting them, maximises intractability in SpyDen, as entries can easily be added or removed from the set of suggested spine centres. Additionally, the faster R-CNN architecture provides a confidence value for each identified spine that can be used to filter spines. Therefore, we have provided a slider in SpyDen that determines the minimum confidence the network must have for a given spine suggestion. The final detection can thus be a mix of automatic and manual spine locations to achieve the best result, as seen in Fig. 2B.

Throughout the development of SpyDen, we trained the underlying neural network on a wide variety of experimental image types, thus ensuring robustness across a wide variety of experiments (for details of the various experimental images used, see Table S4). Nonetheless, we acknowledge that our network may not perform at the desired level for all images and note that this is still an open challenge in the field. However, inspired by the approach of DeepD3 (Fernholz et al., 2023), we include the ability not to use the default SpyDen network and instead provide a self-trained one that takes a 2D image as its input and generates a list of locations as its output (confidences can be arbitrarily set). By pressing the “Set NN (default)” button (see box ii of Fig. S1), SpyDen includes the option to download SpyDen’s default network or provides the path to a custom implementation. We emphasise that the neural network approach is optional for analysing the spines, and the fully manual procedure can also be utilised.

#### Automatic ROI generation

As the above choice of spine identification algorithm only provides the spine centres, SpyDen necessarily includes a separate algorithm for spine segmentation. This algorithm takes the spine locations and then generates ROIs that can be used to analyse those spines.

Our approach relies on the inherent features of the spines to generate a polygon with editable vertices and provides three distinct advantages over common neural network-based approaches:

1. Using our approach leads to an ROI that consists of a polygon with editable vertices. Each vertex can be moved or deleted, and new vertices can be added to enhance the ROI, allowing for significant enhancement of the results once the ROIs are generated. The ROI generation algorithm parameters can also be adjusted to provide better automatic results.
2. By using a heuristic approach and relying on the spine features themselves, the algorithm is largely independent of image quality (unlike pre-trained neural network approaches), leading to a fundamentally robust algorithm across different experimental paradigms.
3. The rules that define the ROI generation are intuitive and can be tweaked, unlike the black-box approach of a pure neural network approach. Coupled with the open-source nature, this allows for altering the ROI generation algorithm of SpyDen as part of our inherent design goals.

The ROI generation then proceeds: a point interior to the spine, *s*_0_ = *(x*_0_*, y*_0_), is supplied via the manual or automatic procedure. We then generate eight outward-pointing rays representing the cardinal and ordinal directions (N, S, E, W, and NW, SW, SE, and NE, respectively). Mathematically, we represent the *i*^th^ point along each of these rays as

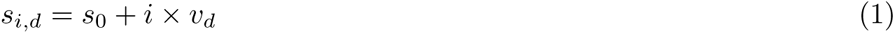

where *v_d_* is a given direction. An example of these rays can be seen in the red arrows of Fig. 2C. For each ray, we additionally introduce a counter *c_d_*. This counter tracks how many times certain rules related to the spine morphology are broken as we step along the ray. When *c_d_* exceeds a certain value *n*, we halt the progression in this direction and denote this point as the edge of our ROI. The counter *c_d_* is independent for each ray, so the ROI generation will progress as long as one of the rays continues to increase.

Examples of the rules we use in SpyDen include the *luminosity drop-off rule* (when the value of the luminosity at point *s_i,d_* falls below a factor of the initial luminosity, *l*_0_) or the *other spine rule* (when the point *s_i,d_* is closer to the centre of another spine than to its own spine centre). Examples of the locations where these rules are broken can be found plotted in different colours in Fig. 2C. The exact details of these rules can also be found in the supplemental material.

Once the counters of all the rays have exceeded the value *n*, i.e., enough rules have been broken for all directions, we generate an octagonal shape encompassing the chosen spine, *s*_0_, where the vertices are given by the final points along each ray. To increase the robustness of the algorithm, we then repeat this process by generating four additional polygons where the initial point is perturbed by one pixel in the cardinal directions. The resulting vertices of the five polygons are then averaged to generate a final polygon that has decreased dependence on the exact location of the initially provided spine position. The exact details of this algorithm can be seen in the supplemental material and Algorithm 3.

Finally, we comment on the difference between the ROI generation if *luminosity mode* or *area mode* is selected. Both these quantities are reliable proxies for the strength of a synaptic connection and have been used extensively to study synaptic plasticity (Matsuzaki et al., 2004; Hayashi-Takagi et al., 2015; Eggl et al., 2023; Chater et al., 2022). Nonetheless, care must be taken when generating the ROIs to study these quantities with temporal components. When studying the luminosity, generating an ROI encompassing the maximum extent of the spine over the entire period is critical. This allows for accurate measurement of the spine dynamics as the luminosity changes within the ROI. On the other hand, when studying the area, we need to generate an ROI that only encompasses the spine at *that* timepoint. Then, the areas of those ROIs dynamically change as the spine shrinks and grows. Therefore, in SpyDen, there is only one ROI in *luminosity mode* and an ROI for each snapshot in *area mode*. Please see tables S2, S3 for a detailed description of the output generated by the synaptic analysis in *luminosity mode* and *area mode*, respectively.

### Motion correction

SpyDen also includes the capability of studying the temporal dynamics of dendrites and their spines. However, given that biological structures inevitably shift in time, be it through their innate motion or the noise of the experimental paradigm, it is critical to introduce a process to correct these shifts. Additionally, this movement may occur globally when the shifting is uniform for the entire image or on a local level, where individual sections may move independently from the rest of the image.

To account for these effects and accurately track the ROIs in time, we employ a phase cross-correlation approach (Foroosh et al., 2002; Guizar-Sicairos et al., 2008). Unlike spatial-domain algorithms, phase cross-correlation relies on the frequency representations of the images and, thus, is more robust to noise and occlusions. Furthermore, in comparison to directly calculating the cross-correlation of the two images, the spectral approach is significantly faster to calculate. This approach is particularly attractive because it can be applied to images of any size without significantly increasing computational time and can be used for global and local motion correction. In SpyDen, we find the necessary translations to account for the global shift and then introduce individual shift corrections via the same algorithm for each individual ROI. This algorithm is implemented by using skimage (Van der Walt et al., 2014) to correct for both dendritic and synaptic shifting.

### Puncta analysis

Localising and calculating the size of fluorescent bright puncta is a crucial component in the analysis performed on images following hybridisation techniques such as fluorescent *in situ* hybridisation (FISH) or single molecule FISH (smFISH). Such techniques result in a fluorescent bright puncta-like signal with a dark background (for example, see Fig. 2E). Thus, given a dataset of multi-channel and/or multi-time images, SpyDen detects puncta in each of the provided images. This is achieved by using the Laplacian of Gaussian (LoG) technique (skimage implementation) to automatically detect fluorescent spots in the experimental images. This method requires a threshold value, defined as the absolute lower bound for scale space maxima. Nevertheless, the optimal threshold value may differ for distinct neurological structures. As SpyDen provides the ability to analyse fundamentally different biological structures (for example, spines, dendrites, somata, etc.), we provide two sliders, labelled “Threshold dendrite” and “Threshold synapse/soma”, respectively, to set the threshold values for each of the two types of ROIs independently. For example, given the image *I*(*t, c, x, y*) and an ROI, *R*(*t, c, x, y*), in image *I*, the threshold, *t_R_*(*t, c*), for *R* is determined as follows:

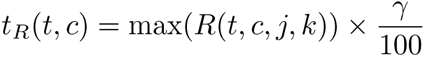

Here *R*(*t, c, j, k*) is the array containing fluorescent intensity value ∀*j, k* ∈ ROI*_I_* in channel *c* and at time *t* and *γ* represent the values of the time and channel sliders. Additionally, SpyDen includes the ability to detect puncta of various sizes by changing the minimum and maximum standard deviation of the Gaussian kernel using the provided puncta size slider. Finally, we omit puncta detection in ROIs whose intensity is below the background noise level to improve the efficiency of the algorithm and to avoid detecting erroneous puncta.

## Performance evaluation

As SpyDen consists of three fundamental analysis components (dendritic, synaptic, and puncta analysis), we perform three separate analyses to determine the effectiveness of using this tool. To this end, we take a variety of datasets that represent realistic experimental use-cases.

### Dendritic analysis

The synaptic and dendritic algorithms are evaluated on three distinct datasets to enhance the validity of the verification and prove the robustness of SpyDen to different experimental paradigms. These three datasets, publicly available and already published, are the Chater (Chater et al., 2022), Helm (Helm et al., 2021), and Cultured datasets (the last of which is unpublished and provided as part of this work). The first dataset contains examples of neurons from organotypic slice culture, while the latter two contain neurons from dissociated cultures, which makes the combination an attractive set of data for the verification of SpyDen. One key distinction between the datasets is the different experimental resolutions, a feature that many neural networks rely on to achieve their performance. Additionally, the difference in contrast between background noise and foreground means that conventional methods may struggle. For example, images from each of these datasets, see A, B, and C of Fig. 3.

**Figure 3:**
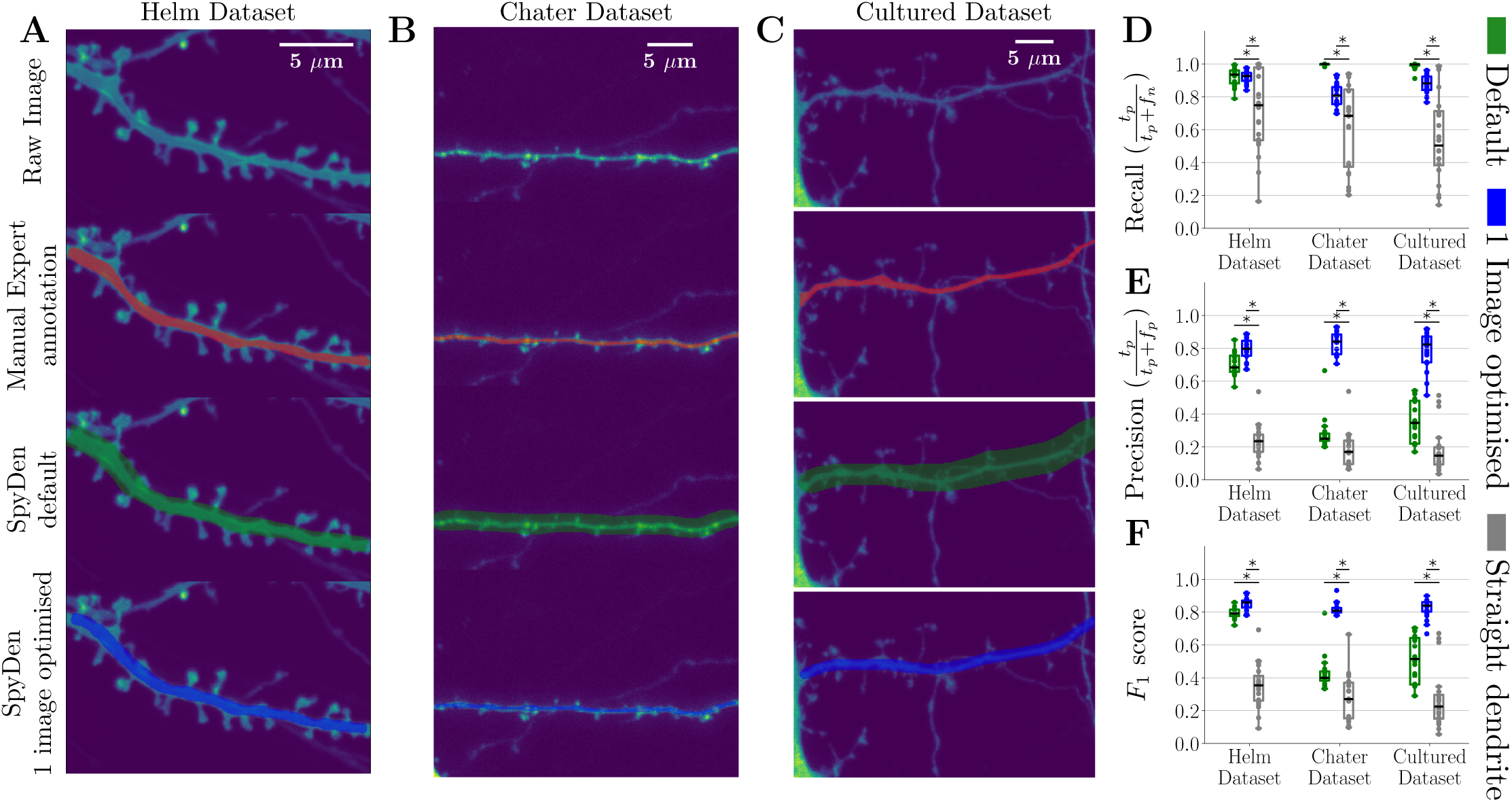
The dendritic segmentation in two different datasets reliably obtains performance comparable with manual expert evaluation. **A)** Example dendrite segmentation of a dendrite from the Helm dataset (Helm et al., 2021). From top to bottom: Raw image, showing the dendritic stretch and associated spiny protrusions, manual annotation of of the dendrite by an expert overlaid in red, segmentation achieved with the SpyDen tool if default values are used in green and optimal segmentation if the algorithm parameters are augmented (blue). **B)** As in A), but using a set of data from the Chater dataset (Chater et al., 2022). Colour scheme remains the same. **C)** As in A), but using a set of data from the Cultured dataset. Colour scheme remains the same. **D)-F)** To measure the performance of dendritic segmentation, we use calculate the recall (*D*) (true positive results divided by the number of all samples that should have been identified as positive), precision (*E*) (number of true positive results divided by the number of all positive results) and *F*_1_-score (*F*) which is a measure of our algorithms accuracy. In each figure, the green and blue refer to the performance of the SpyDen default and adapted parameters, respectively. Additionally, we calculated a baseline (called the *straight dendrite*), where a diagonal chord (2*µm* width) is drawn between the dendritic start and end-points and designated as the dendrite (grey). *^∗^* refers to *p* < 0.05.

We begin by studying the performance of the dendritic segmentation algorithm and compare this to the results of a manual expert evaluator. Each dataset comprises 20 dendrites. Natural use of SpyDen includes directly applying the results arising from the automatic algorithm, adjusting that result by modifying the medial axis path of the dendrite or by altering the algorithm parameter sliders. To mimic this natural use, we compare the expert to two distinct segmentations:

- Default: the automatic segmentation when the default parameters of SpyDen were used.
- 1 image optimised: the automatic segmentation resulting by altering the algorithm sliders akin to natural use for one image of the new dataset. These parameters are then used for all other images of that dataset

Examples of dendrites from each dataset and the three different segmentations can be seen in Fig. 3 in green and blue, respectively.

Using the calculated segmentations, we can then generate metrics that quantify the performance of the SpyDen algorithm when compared to the ground truth (expert annotation). In our case, the segmentation task is equivalent to a classification task, as we are trying to classify each pixel as part of the dendrite. Additionally, this is an imbalanced task, as there are many more non-dendrite pixels than dendrite pixels. Metrics commonly used to evaluate the performance of classification algorithms in imbalanced datasets are the recall, precision, and *F*_1_ scores (Powers, 2020). Recall measures the ability of a model to correctly identify all relevant instances of the positive class (true positives) out of the total actual positive instances, precision measures the ability of a model to correctly identify positive instances without including too many false positives, and the *F*_1_ Score is the harmonic mean of precision and recall, defined as

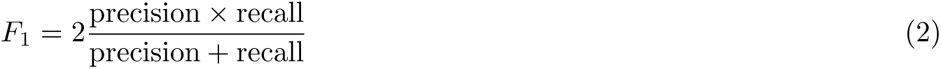

and provides a trade-off between these two metrics.

The results calculated for the two SpyDen segmentations can be seen across the Helm, Chater, and Cultured datasets in Fig. 3D-F. These figures also include the full equations of recall and precision. As each dataset has different characteristics, we expect to see variations in these metrics as the performance of SpyDen will naturally fluctuate. The algorithm annotations of the Helm dataset have high recall (≈ 0.9) and precision for both the default and altered parameters (≈ 0.8). This is at a level where there is little discernible difference between these two segmentations. We attribute this to the fact that the parameters of the dendrite algorithm were developed and tested with the experimental features of this dataset in mind. Therefore, altering the algorithm parameters to obtain a better segmentation leads to only a small improvement in the classification metrics. This leads to an average F1 score of 0.85, indicative of a well-performing model with a good balance between minimising false positives and false negatives.

To further understand the performance of our model, we design a naive segmentation strategy that conceivably could generate reasonable dendritic segments, which we call the “straight dendrite” (see grey lines points in Fig. 3D-F). This naive strategy is defined by taking the provided start and end points and drawing a chord of a certain thickness between these points. We found that the best average *F*_1_ metric was achieved when this thickness was ≈ 2*µm*, which aligns with experimentally reported dendritic widths (Stuart et al., 2016). Nevertheless, this baseline performs significantly lower than both the SpyDen segmentations and only achieves an average *F*_1_ value of 0.4 and 0.3 for the Helm and Chater datasets, respectively. As the dendrites are not perfectly straight segments, this strategy is unable to deal with dendrites that may curve and thus deviate from the chord. This leads to low precision and highly variable recall.

Turning to the Chater dataset, we see that the default SpyDen parameters lead to a worse performance, which can be primarily attributed to the lower contrast between the background and the dendrite. Thus, the default parameters (particularly the variance of the Gaussian filter used to detect the edges of the dendrite) tend to overestimate the size of the dendrite, leading to a high recall rate (all points in the dendrite covered). However, this leads to a low precision rate as the segmentation includes many points of the background that are not part of the dendrite. Nonetheless, we still outperform the baseline strategy. With minimal adjustments to the algorithm parameters, we can achieve a performance comparable to that of the Helm Dataset (*F*_1_ score 0.82). As SpyDen algorithm variables can be saved and transferred to other experimental images, the alteration of these parameters was only performed on one Chater dataset image and then applied to all other images. Thus, the dendrite segmentation of SpyDen can reliably obtain a satisfactory result across a wide array of experimental conditions. We note that the results of the Cultured dataset mirror those of the Chater dataset, i.e., the default SpyDen values represent an overestimation of the dendritic stretch, with high recall but low precision. Nonetheless, by augmenting the parameters to correct for this over-estimation, we achieve a reasonable result with an *F*_1_ score of 0.85). We note that with both approaches using SpyDen (automatic and augmented), the F1 performance is significantly better than the baseline.

We highlight that augmented segmentation was achieved without altering the path of the dendritic stretch (via the editable nodes). Therefore, an even higher *F*_1_ score is possible. Nevertheless, as we aimed to mimic fast and efficient natural user analysis, we restricted ourselves to only changing the transferable parameters between different experimental images.

### Spines

We now turn to verifying the abilities of SpyDen to accurately measure dendritic spines, using the previously employed datasets as they image both dendrites and spines. The “ground-truth” ROIs are obtained using manual annotation of the spines by expert evaluators, which are then compared against the results from SpyDen. As mentioned by Fernholz et al. (2023), using human expert annotators as the ground truth can be problematic as biases due to experience and experimental expertise can lead to different results. This means that comparing to only one annotator could lead to better or worse performance based on the results of that expert. To combat this, we used a set of three expert annotators that each worked on different datasets to hopefully provide the most “accurate” ground truth.

We begin by comparing the average SpyDen ROI luminosities against the ground truth luminosity as evaluated manually. The luminosity is a measure of the spine head volume which is a proxy for synaptic strength (i.e., the number of AMPA receptors) and is used in a wide variety of studies to understand synaptic plasticity (Matsuzaki et al., 2004; Hayashi-Takagi et al., 2015; Eggl et al., 2023; Chater et al., 2022). Here, the SpyDen ROIs are generated using the 1-image optimisation approach mentioned above, i.e., we only change the parameters of the automatic algorithms once for each dataset and then leave them untouched. Nonetheless, we note that the results presented here could be further improved by taking advantage of SpyDen’s capability to change the ROIs directly.

We find that the luminosities of the spines, as calculated by SpyDen, are in good agreement with those generated by the expert annotator, as can be seen in Fig. 4A). When pooling all three datasets, the correlation exceeds 0.9, and the slope of the linear relationship between is ≈ 1. Additionally, we maintain this strong correlation when studying each dataset individually, albeit with slightly different linear relationships (see Fig. S6 for individual linear fits). Synapses are also dynamic structures that change constantly, a process known as synaptic plasticity. Therefore, we additionally need to verify that the algorithms underlying SpyDen can track these structures and provide accurate luminosity values over time. To this end, we took advantage of the temporal data provided by the Chater dataset and applied the automatic spine tracking provided by SpyDen. The results were then compared to those from an expert annotator (see Fig. 4B, where the blue, red and purple colours refer to three different analysed dendrites). We note that a good agreement is found between the two evaluations (r=0.79). This good performance was achieved without any manual intervention to further enhance the results, which can be easily performed in SpyDen.

**Figure 4:**
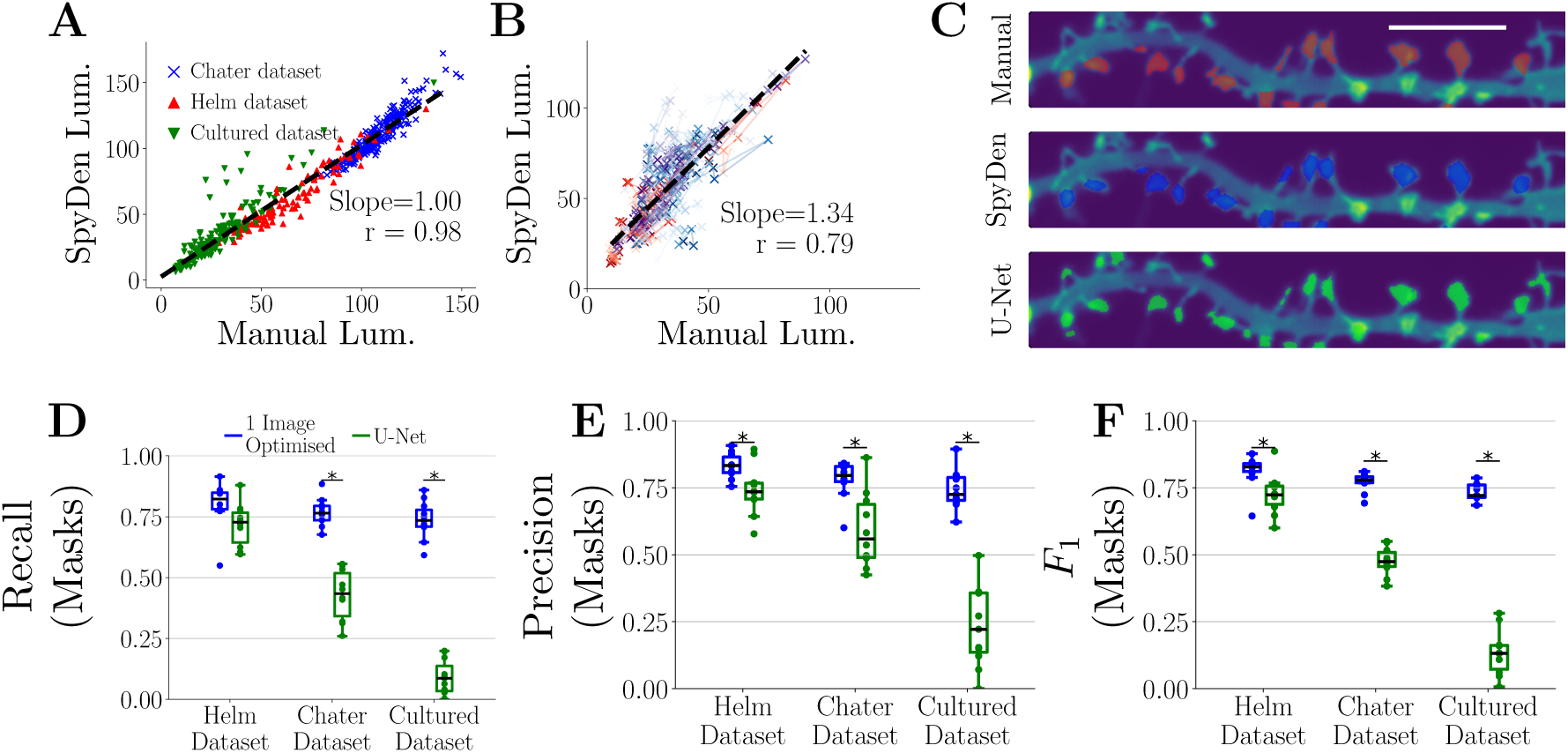
The synaptic detection and ROI generation of SpyDen achieves reliable performance when compared to a U-Net approach. **A)** To verify the results of the SpyDen analysis pipeline, we compare the average luminosity of the SpyDen ROIs against expert manual evaluation. The ROIs of SpyDen can be used to calculate the luminosity of each spine and compared against the manually evaluated luminosity. We achieve robust agreement with the expert evaluation (for the three datasets), as can be seen with the overlaid linear fit. **B)** Using temporal data from the Chater dataset allows us to determine the spine tracking capabilities of SpyDen. We note that a good agreement is achieved between the luminosity calculated by SpyDen and that of the manual annotation of the expert. **C)** Example ROIs generated manually by the expert (red), the heuristic SpyDen rules (blue) and a U-Net architecture (green). The white bar refers to 5 *µm*. **D)-E)** We can directly compare these ROI masks to obtain further verification of SpyDen’s capability. From left to right, we present the recall, precision and *F*_1_ arising from SpyDen (blue) and the U-Net (green) while using the manual annotation as the ground truth. *^∗^* refers to *p* < 0.05.

Next, we compare the ROI masks directly with those generated by the expert (as the luminosity can be obtained even without accurately covering the spine). Additionally, to see how our technique performs compared to other approaches, we trained a U-Net on the Helm dataset to output spine ROIs (Ronneberger et al., 2015). The U-Net takes the entire image as input and provides a binary mask of the same dimensions, highlighting possible spine areas. Given this structure, the U-Net might mark spines not part of the area we wish to analyse (e.g., lying on another dendritic branch). These extra selections would then unfairly decrease the accuracy of the U-Net, so all masks that the U-Net provided that lay outside the analysis region were removed. Nonetheless, we note that generating these extra (possibly erroneous) ROIs may be a problem that real users can encounter in real-world settings and that SpyDen does not have. Additionally, we note that the performance we would get by training the U-Net on all three of our test datasets would lead to trivially good results. However, training the network on this larger data set is computationally intense and not feasible for most end-users. Therefore, to mimic likely real-life analysis, we take the network and only train it on one dataset (in this case, the Helm dataset) and then apply it to all three datasets.

SpyDen, on the other hand, is not “trained” on a given dataset or resolution and instead consists of deterministic algorithms that rely on the biological structure of the spines and, therefore, can generalise beyond the datasets we have used here. Additional generalisation is achieved by enhancing the automatic results by altering the parameters underlying the algorithm and then applying these to all other images of the same experiment. This requires no knowledge of coding or computational resources, unlike retraining the neural network using the U-Net approach. We believe the above U-Net implementation represents the fairest comparison for SpyDen.

To evaluate the performance of the SpyDen and U-Net ROIs, we use the previously defined recall, precision, and *F*_1_ metrics, using the expert annotation as the “ground truth” baseline (Fig. 4D-F). Examples of these three ROIs can be seen in Fig. 4C, where red, green, and blue refer to the manual, SpyDen, and U-Net implementations, respectively. As expected, the U-Net and SpyDen perform at similar levels for the Helm dataset. However, SpyDen performs significantly better than the U-Net for the other two datasets, which follows intuitively as the U-Net is unable to adapt to the new experimental conditions. SpyDen achieves ≈ 0.8 for the three datasets, which is indicative of a well-performing model that does not rely on the image’s resolution to generate accurate results, making it a promising tool for evaluating spines across a wide array of datasets.

We also evaluated the performance of the SpyDen neural network in identifying spine centres in the three datasets (see Fig. S4I). We note that we obtain a consistent *F*_1_ performance of ≈ 0.7 but emphasise that the spine centres provided by the neural network can be enhanced simply by adding or removing the suggested points with a single click, which can not so simply done in the U-Net architecture.

### Puncta

To test the accuracy and efficiency of the SpyDen puncta analysis, we compared the results of SpyDen on two recently published data sets (Ciolli Mattioli et al., 2019; Fonkeu et al., 2019). The first dataset consists of images of mouse primary cortical neurons, in which two different Cdc42 isoforms were labelled using single molecule mRNA FISH (shown in Fig. 5A). Dataset2 contains images of rat dissociated hippocampal cultured neurons, on which mRNA FISH was performed for CaMKII*α* (shown in Fig. 5C, left). In the original work, quantification of smFISH puncta for both of Cdc42 isoforms (namely, E6 and E7) was performed using StarSearch (RajLab), which is a well known and frequently used tool for smFISH puncta quantification. The results from SpyDen’s analysis were comparable to those from the original work in terms of the percentage localisation of the two isoforms between soma and neurites (see Fig. 5D, E). Similarly, in the original work, dataset 2 was analysed using Neurobits (Tushev et al., 2018). Again, the quantification of CaMKII*α* mRNA from SpyDen in somatic and dendritic compartments matched well with the results obtained using *Neurobits* (see Fig. 5F). Furthermore, we obtained comparable results in terms of the count of puncta detected in each compartment (soma vs neurites) in both data sets (see Fig. 5G, H, I and Fig. S7). One of the advantages of using SpyDen puncta detection is that it allows manual enhancements by providing the option to inspect and alter the underlying algorithm parameters (such as threshold values or puncta size). Additionally, SpyDen’s approach promotes efficient evaluation of the puncta; instead of identifying puncta in the whole image (as is done in *Neurobits*), SpyDen looks for puncta only in designated ROIs. This greatly reduces the time spent analysing each image. Another advantage of using the SpyDen puncta detection is that we provide video tutorials demonstrating how to use the analysis pipeline. This means that learning how to use SpyDen is much easier than learning how to use other tools that only provide text-based documentation. Moreover, existing tools often do not provide useful statistics which are automatically output by SpyDen, such as the size of each punctum, their location, and minimum and maximum intensity. For a complete list of puncta statistics, see Table S5.

**Figure 5:**
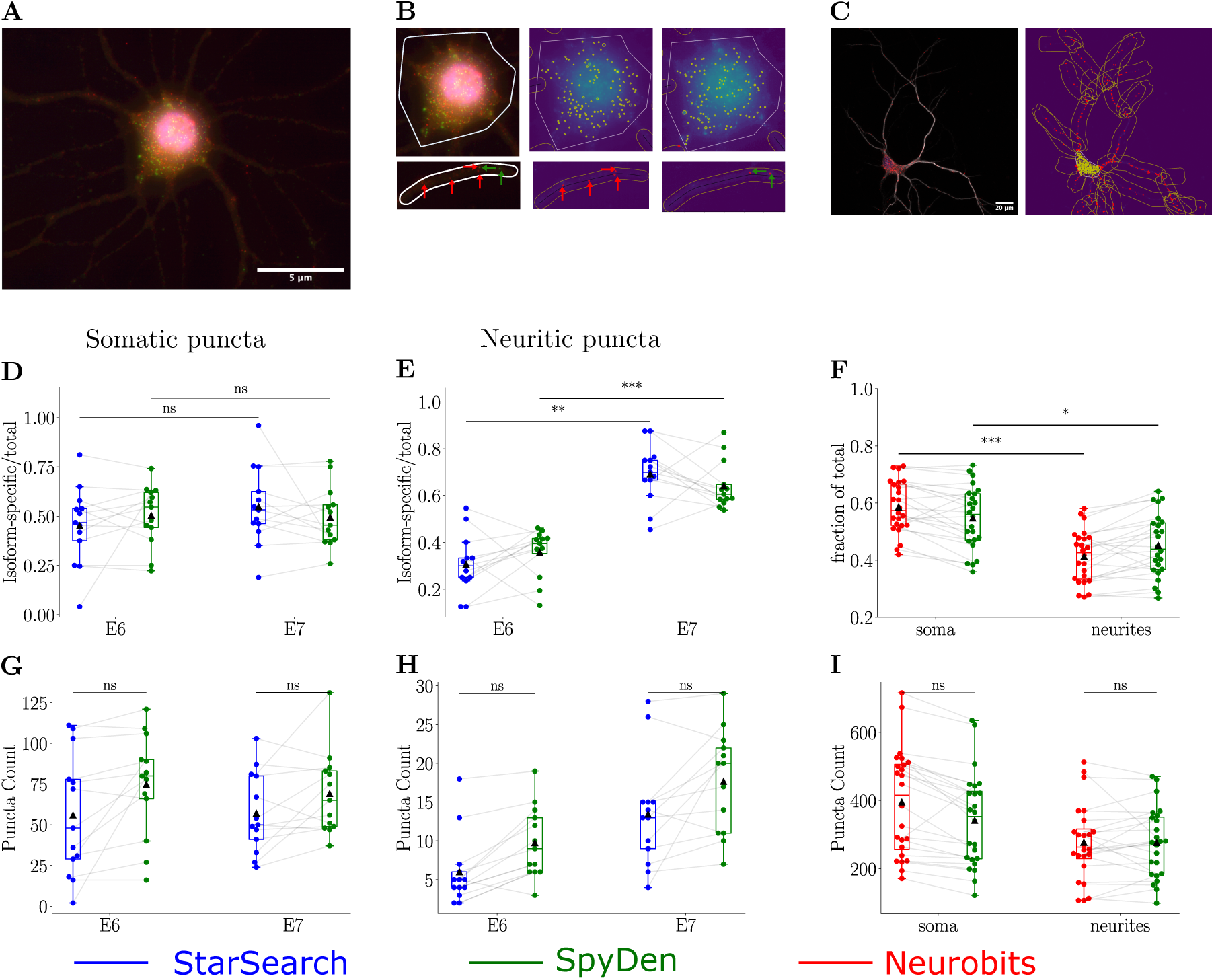
Validation of SpyDen’s puncta detection pipeline on published datasets. **A)** smFISH of Cddc42E6 (green channel) and Cdc42E7 isoforms (red channel), cell body is labelled with DAPI (in blue channel). Arrows point to individual punctum. **B)** Zoomed-in image of segmented soma and an example dendrite with smFISH puncta for both isofroms (E6 in green,E7 in red) on left and the same pucta detected by SpyDen on right. **C)** FISH of CaMKII*α* mRNA in red with fluorescent immunolabeling of MAP2 for neurites (on left) and corrosponding somatic, dendritic and identified mRNA puncta by SpyDen on the right. **D)** the ratio of isoform specific to total Cdc42 in somata based on puncta count by StarSearch and SpyDen. Ratios based on results from SpyDen is comparable to that of StarSearch. **E.** Same as in D but for neurites. Isoform E7 has statistically higher localisation that E6. The same is obtained from the results of SpyDen analysis (*: p-value < 0.05,**: p-value < 0.01, ***: p-value < 0.001). **F.** Soma vs neurites fraction of total CaMKII*α* mRNA with higher localisation in soma. The analysis using SpyDen is able to replicate the results obtained using Neurobits. **G-I** Comparison of smFISH puncta count in by StarSearch vs SpyDen in soma (in **G**), in neurites (in **H**) and FISH puncta count by Neurobits vs SpyDen (in **I**)

## Conclusion

With SpyDen, we have presented a robust and versatile tool for studying continuous and discrete molecular localisations inside dendrites and spine compartments. While we developed SpyDen with dendritic compartments in mind, the algorithms underlying SpyDen are also readily applicable to other cellular compartments, e.g. fluorescently labelled nuclei within an area of interest or the number of molecules of interest inside a specific cellular compartment. To aid the adaptation and ease of use, we have incorporated into SpyDen three fundamental principles: *i)* easy-to-use and intuitive operation without prior knowledge of programming due to a graphical user interface and video tutorials, *ii)* it is fully open-source and open-access, containing no proprietary software *iii)* results arising from the ANN-based tracing and ROI algorithms are editable.

Finally, we cross-validated and bench-marked SpyDen performance across different experimental datasets and have shown that SpyDen achieves reliable results when tasked with tracing dendrites, identifying spines, and fluorescent puncta and its performance is on par with expert manual annotators. On top of the automatic analysis performed by built-in algorithms, it can enhance the results, leading to even more reliable analysis. Each step of the SpyDen analysis pipeline can be user-adapted, and the parameters underlying the algorithms can be changed to obtain better performance. Thus, SpyDen provides a baseline performance that increases reproducibility and allows additional customisation by adapting the algorithmically-suggested result. Thus, we believe that the SpyDen framework provides a powerful platform for cellular studies to analyse large neuronal images more efficiently and reproducibly and gain insight into the processes underlying dendrite and synapse dynamics.

## Acknowledgement

We thank all members of the Tchumatchenko lab in particular K. Mousaei, C. Bergman and J. Petkovic for fruitful discussions. We acknowledge support from the University of Bonn Medical Center (S.W.,T.T.), the University of Mainz Medical Center (S.W.,T.T.) and the Joachim Herz Foundation (M.F.E). This project has received funding from the European Research Council (ERC) under the European Union’s Horizon 2020 research and innovation program (‘MolDynForSyn,’ grant agreement No. 945700).

## Data and code availability

Data and code to generate the figures found in this manuscript can be found in the following public github repository Spyden-PaperFigs. The full code base for SpyDen, including instructions on installation and use, can be found at the github repository SpyDen while the compiled executables can be found at gin.g-node.org/CompNeuroNetworks/SpyDenTrainedNetwork.

## Statistical methods

Comparisons between expert manual and SpyDen results were performed using a Studentized t-test and multiple comparisons were corrected for using a Bonferroni correction factor.

## Supplemental material

### SpyDen GUI

**Figure S1:**
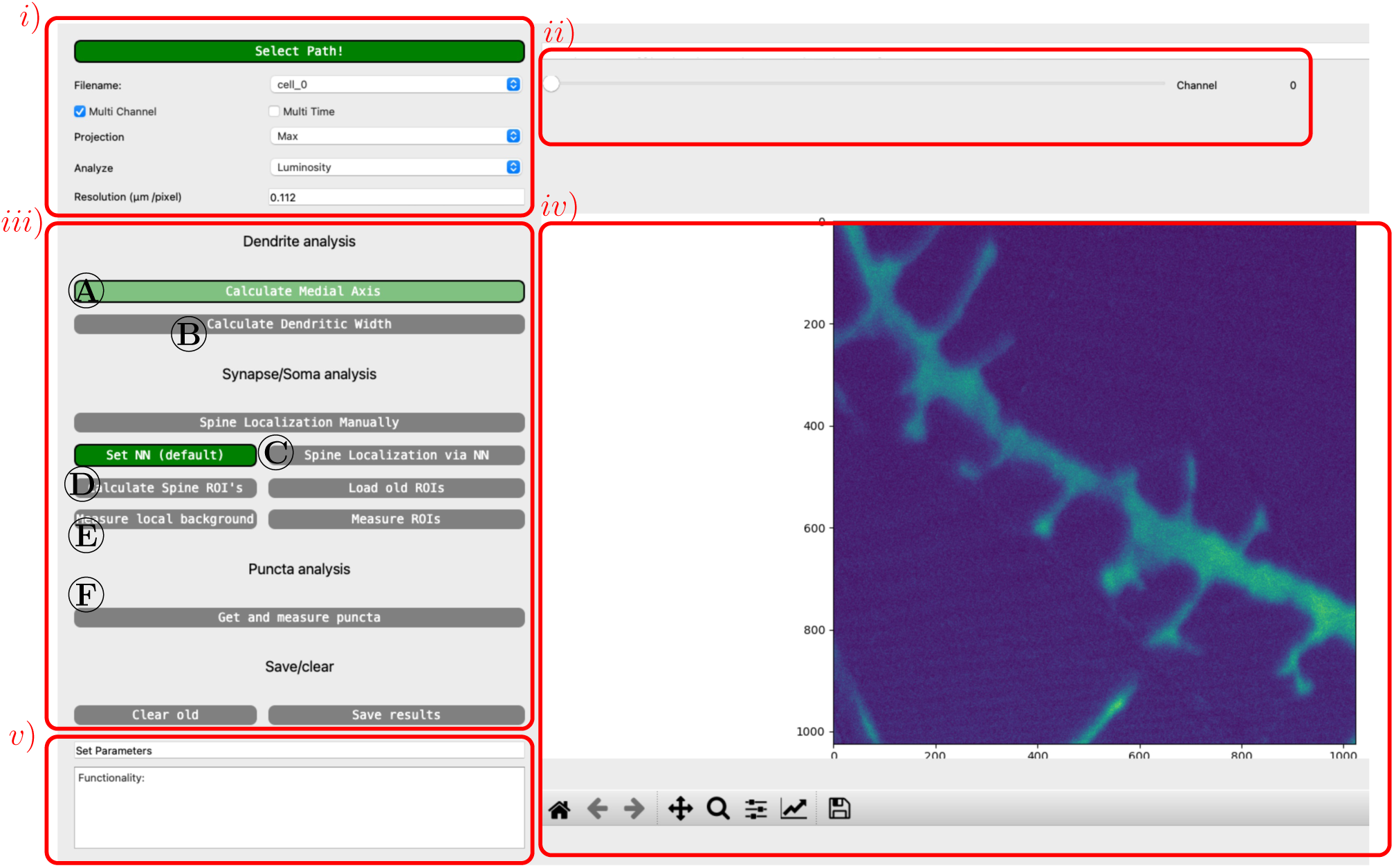
Principle analysis window of SpyDen. We highlight the different components of this window in the red boxes and also show relate the analysis modes seen in Fig. 1 to the buttons that can be selected here.

We emphasise that a full description of the GUI and data analysis pipeline of SpyDen can be seen in a set of video tutorials that we encourage the readers to watch.

The SpyDen GUI is the primary interaction point the user has with SpyDen. Coupled with a suite of video tutorials, SpyDen is easy to pick up and does not assume any programming knowledge to analyse experimental images. The primary analysis window is designed to provide the user with immediate feedback on what they can currently do, as each analysis step is only unlocked once all prerequisites are filled (green for active, grey for inactive). This step-by-step process leads to intuitive analysis, as logical and repeatable steps are performed for every data set.

The different components of the analysis are highlighted with red boxes in Fig. S1. Box i determines the meta-parameters of the data analysis, including the type of z-stack projection, image resolution and whether multiple channels are present. As the algorithms underlying the different analysis pipelines consist of tunable parameters that can affect the final result, we have introduced a set of sliders that appear for the relevant step in box ii. To select the different analysis modes, the user clicks the buttons in box iii. Box iv) provides feedback to the user by depicting the interactable image that can be clicked on, zoomed in on and analysed. Finally, box v is a text box that tells the user the current status of the analysis and what possible keyboard shortcuts they can use.

As the user proceeds through the analysis pipeline provided by SpyDen, the image in box iv changes to reflect the current status of the analysis. Examples of the different states of the image in box iv are depicted in Fig. 1B-G) with the buttons that lead to this behaviour marked in Fig. S1, iii). These include the dendritic analysis (A, B), synaptic selection and segmentation (C-E) and the puncta analysis (F). As the image is built on an existing Python package (matplotlib), all the functionality of saving, panning and zooming are already implemented.

### SpyDen data input

SpyDen takes *.tif*, *.lsm*, *.png* or *.jpeg* files as possible input. This choice of input formats was driven by the fact that the*.tif* format is some of the most widely used image formats in bio-sciences, while *.png* and *.jpeg* files are commonly used for general image files.

SpyDen then provides the option to alter the type of analysis that will be performed, e.g., the type of z-stack projection or whether to include temporal dynamics (to study such dynamics, the different time points need to be provided as separate images). SpyDen additionally requires the input of the experimental image’s resolution (a necessary requirement for some of the subsequent analysis), which, if available, is obtained from the image metadata or provided manually.

### Structure of the SpyDen output

**Table S1:**
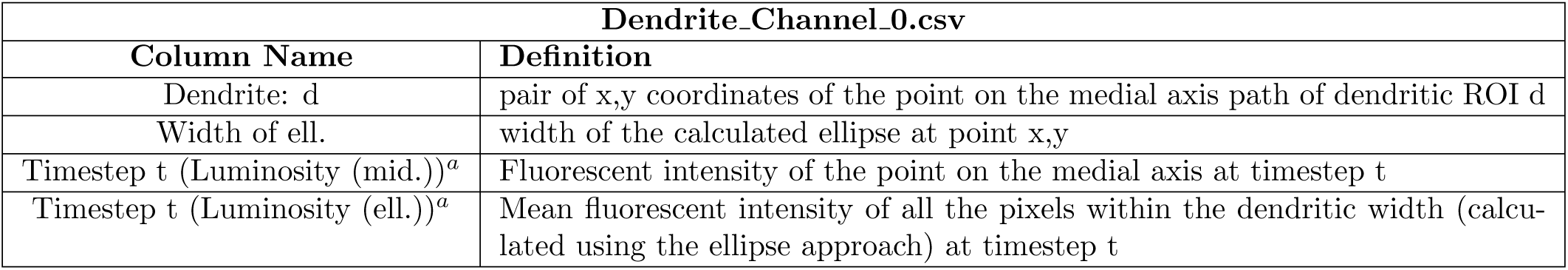
Output file contains measurements for individual dendritic ROIs. *^a^* a separate column is added for each time step. A separate file is created for each channel as well.

**Table S2:**
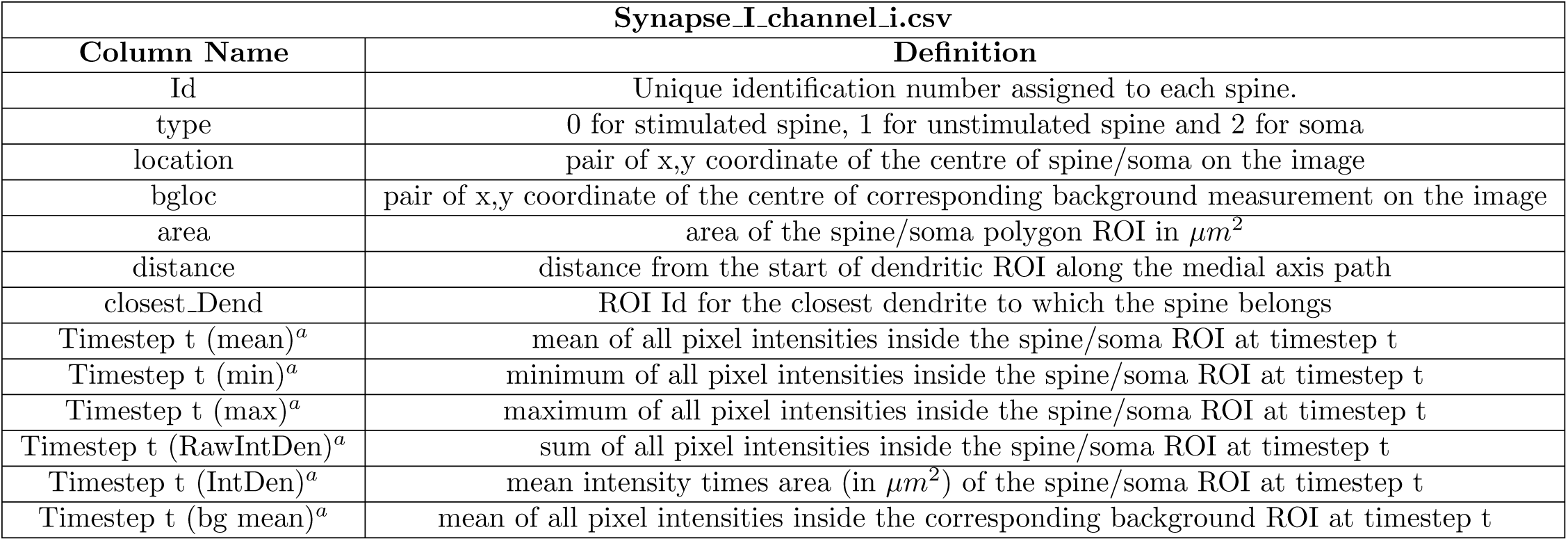
Output file structure generated for spine/soma ROIs when analysed using *luminosity mode*. *^a^* a separate column is added for each time step and one file per channel is created.

| Synapse_I_channel_i.csv |  |
| --- | --- |
| Column Name | Definition |
| Id | Unique identification number assigned to each spine. |
| type | 0 for stimulated spine, 1 for unstimulated spine and 2 for soma |
| location | pair of x,y coordinate of the centre of spine/soma on the image |
| bgloc | pair of x,y coordinate of the centre of corresponding background measurement on the image |
| area | area of the spine/soma polygon ROI in $\mu m^2$ |
| distance | distance from the start of dendritic ROI along the medial axis path |
| closest_Dend | ROI Id for the closest dendrite to which the spine belongs |
| Timestep t (mean)<sup>a</sup> | mean of all pixel intensities inside the spine/soma ROI at timestep t |
| Timestep t (min)<sup>a</sup> | minimum of all pixel intensities inside the spine/soma ROI at timestep t |
| Timestep t (max)<sup>a</sup> | maximum of all pixel intensities inside the spine/soma ROI at timestep t |
| Timestep t (RawIntDen)<sup>a</sup> | sum of all pixel intensities inside the spine/soma ROI at timestep t |
| Timestep t (IntDen)<sup>a</sup> | mean intensity times area (in $\mu m^2$) of the spine/soma ROI at timestep t |
| Timestep t (bg mean)<sup>a</sup> | mean of all pixel intensities inside the corresponding background ROI at timestep t |

**Table S3:** Output file structure generated for spine/soma ROIs when analysed using *area mode*. *^a^* a separate column is added for each time step and one file per channel is created.

| **dend_puncta.csv and soma_puncta.csv** |  |
| --- | --- |
| **Column Name** | **Definition** |
| Id | Unique identification of the spine ROI |
| type | 0 for stimulated spine, 1 for unstimulated spine and 2 for soma |
| location | pair of x,y coordinate of the centre of spine/soma on the image |
| closest_Dend | ROI Id for the closest dendrite to which the spine belongs |
| Timestep t (area)<sup>a</sup> | area the spine/soma ROI at timestep t in unit<sup>2</sup> |
| Timestep t (mean)<sup>a</sup> | mean of all pixel intensities inside the spine/soma ROI at timestep t |
| Timestep t (min)<sup>a</sup> | minimum of all pixel intensities inside the spine/soma ROI at timestep t |
| Timestep t (max)<sup>a</sup> | maximum of all pixel intensities inside the spine/soma ROI at timestep t |
| Timestep t (RawIntDen)<sup>a</sup> | sum of all pixel intensities inside the spine/soma ROI at timestep t |
| Timestep t (IntDen)<sup>a</sup> | mean intensity times area (in $\mu m^2$) of the spine/soma ROI at timestep t |

### Effect of the algorithm parameters on the dendritic segmentation

**Figure S2:**
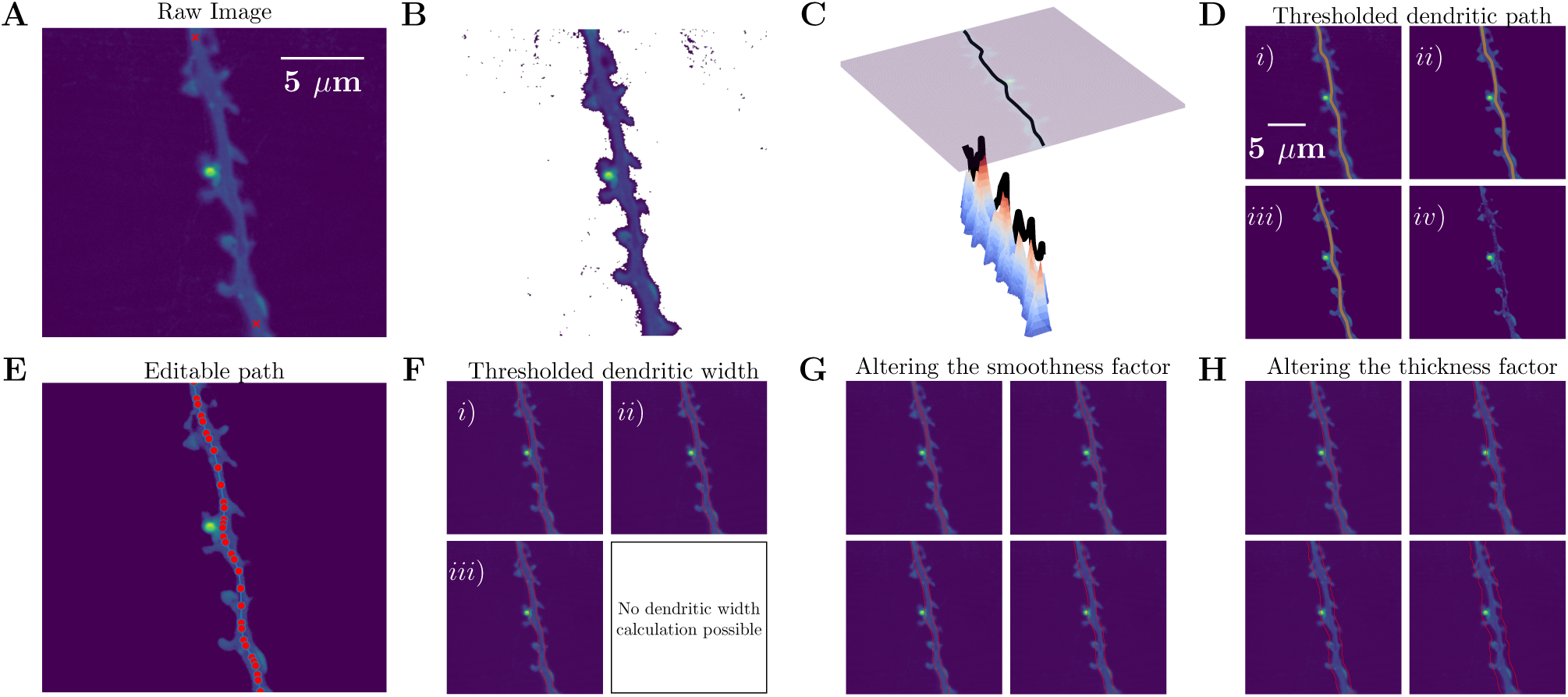
Applying the median threshold to the raw image allows us to generate better medial axis paths for the dendrites and subsequently better approximations of the dendritic width. **A)** Example input image that is provided to the code with start and end points of the dendritic segmented marked in red by the user. **B)** Given this image, SpyDen generates a filtered version of the image, where all noise is removed from the image and only key features are retained. Here, we use the default threshold value. **C)** Given this filtered image SpyDen generates a weighted matrix (see 3D plot) where it finds the shortest path that traverses the highest points. These highest points represent the centre of the dendrite. **D)** By using different threshold values, more or less features are filtered out. This leads to slightly different optimal dendritic paths (marked with the orange line). Each of the 4 sub-images (*i* to *iv*) depicts a slightly higher user-set threshold. We note that case iv) filtered too much of the dendrite out and so no optimal path was found. **E)** Once a suitable path is calculated, the user can make fine corrections to the code by moving editable nodes along the suggested path (highlighted by the red circles). **F)** Using the selected threshold and dendritic path a dendritic width can be calculated (red lines on either side of the dendrite). We note that it was not possible to find the dendritic path and subsequently the dendritic width for case iv) as the threshold eliminated significant parts of the dendrite. Thus, there is no viable path between the start and end points. **G)-H)** A set of tunable parameters allow for user interaction with the dendritic width calculation. These are the smoothness factor in (G), which determines the amount of pixels that are used to calculate the length of the outward pointing normal from the medial axis and the width multiplication factor in (H) which multiplies the outward pointing normals by a set value. Altering these values can have a substantial effect on the calculated width and allow for a significant amount of flexibility for the user. The threshholded dendritic with from *F* iii) was used.

### Neural network approach to spine identification

To train the neural network, three separate datasets were employed. Examples of these images can be seen in Figure S4. More concretely the details of the images can be seen as follows:

**Table S4:** Experimental details of the network images used in the spine detection ANN.

| Dataset | Type of cell | Resolution | Reference | Use in network |
| --- | --- | --- | --- | --- |
| Helm | Hippocampal cultured neurons | $0.02021 \mu m$ | Helm et al. (2021) | Training + Testing |
| Chater | Organotypic hippocampal slice culture neurons | $0.66 \mu m$ | Chater et al. (2022) | Training + Testing |
| Deep3D | Organotypic hippocampal slice culture neurons | $0.094 \mu m - 0.1245 \mu m$ | Fernholz et al. (2023) | Training |
| Cultured | Hippocampal cultured neurons | $0.29 \mu m$ | N/A | Testing |

**Figure S3:**
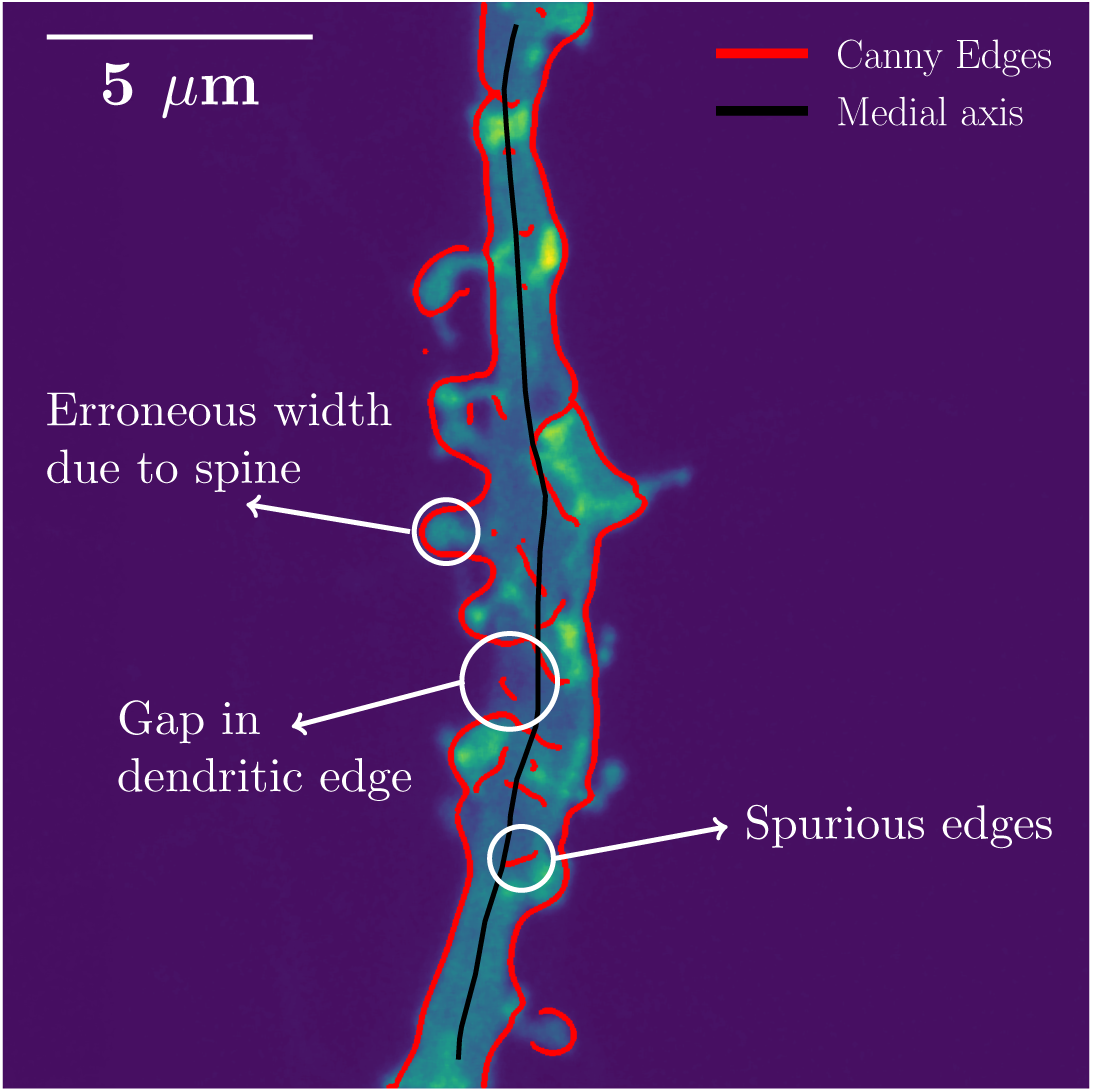
Applying the canny-edge detection algorithm to a filtered version of the experimental images provides an acceptable set of dendritic edges. However, as illustrated by the white circles, several problems preclude using the edges directly. Instead, we apply the ellipse approach seen in Fig. 2A-E) to enhance the dendritic segmentation.

**Figure S4:**
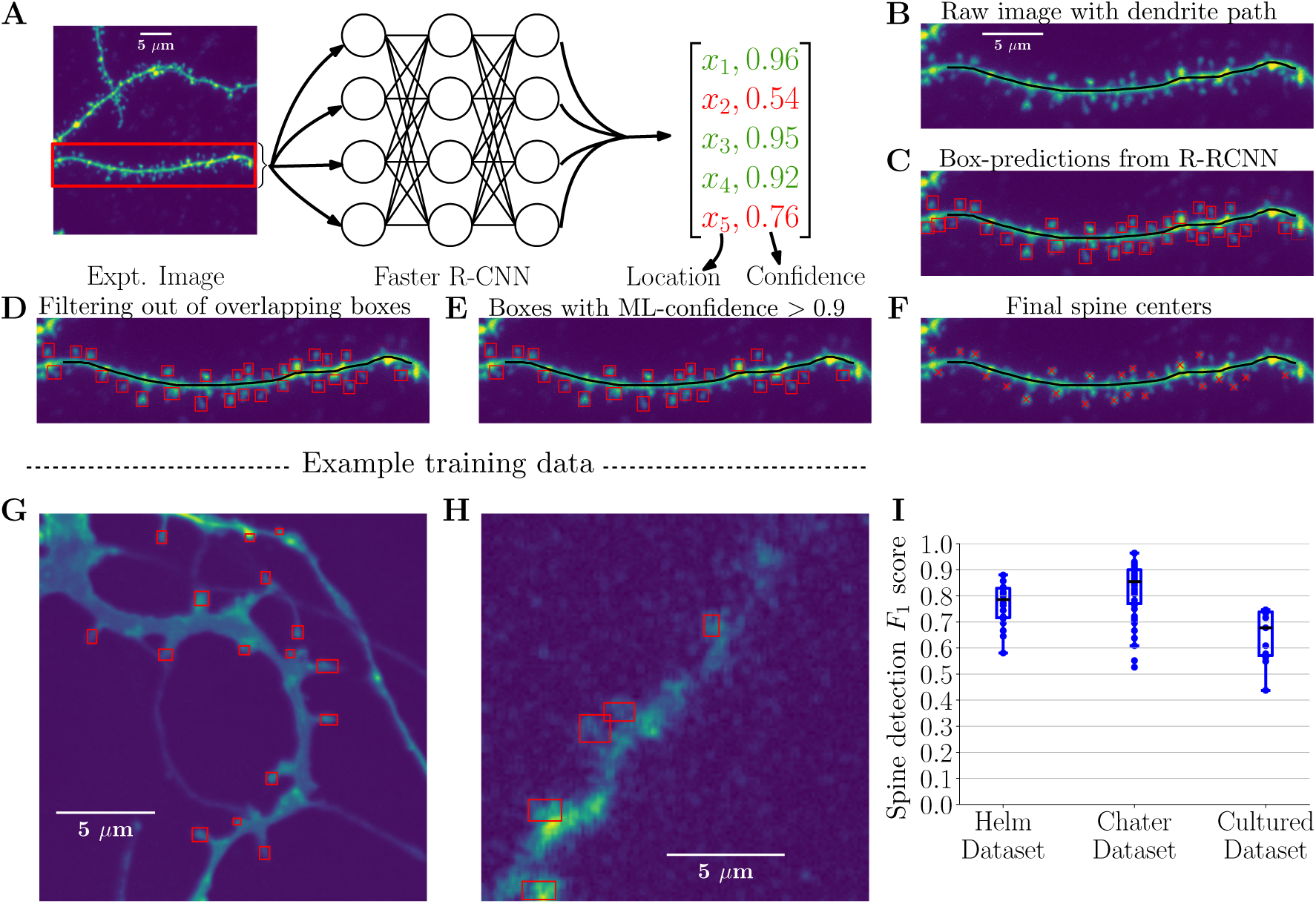
Depiction of the neural network approach with example images used in the spine detection in SpyDen. **A)** Schematic depicting a simplified view of the neural network that is described in (*C* -*F*). Initially, a bounding box is drawn around the dendritic stretch of interest (also seen in *C*). Then, this information is fed to a pre-trained neural network which outputs a list of bounding boxes within that dendritic and the associated confidence of the neural network. The user can then choose to filter the output of the network based on the confidence. Here, we have set the confidence threshold at 0.9, so the suggested points *x*_2_ and *x*_5_ are excluded from further analysis. **B)** Example input that is fed into the neural network. Depicted in black is the medial axis path calculated in the tool as part of the previous step of the analysis. Using this image, the pre-trained neural network (based on a Faster R-CNN architecture (Girshick, 2015), but modified for our purposes), we are then provided a set of bounding boxes and confidence scores for each proposed identified spine. The opacity of each box is defined by the confidence of the algorithm. **D)** When the algorithm suggests multiple overlapping bounding for the same spine, only the bounding box with the highest confidence is selected. **E)** As part of the neural network approach, the user can filter out bounding boxes below a certain confidence score. Here, we have taken the boxes from (c) and filtered out those that have a confidence level less than 0.9. **F)** Finally, using the bounding boxes calculated in (d), we can generate the suggested spine centres (marked with a red cross). These points are then used to generate the bounding ROIs for the subsequent analysis steps. **G, H)** Example images (with manually generated bounding boxes) taken from the two additional datasets used to train the SpyDen spine detection network. These datasets are the Helm dataset (Helm et al., 2021) and Deep3D (Fernholz et al., 2023) datasets. **I)** Evaluation of the spine centre detection of the SpyDen neural network using the *F*_1_ metric. We note that the automatic procedure to generate spine centres (and which can be augmented with manual selection/deletion) leads to reasonable results across the three different test datasets.

**Figure S5:**
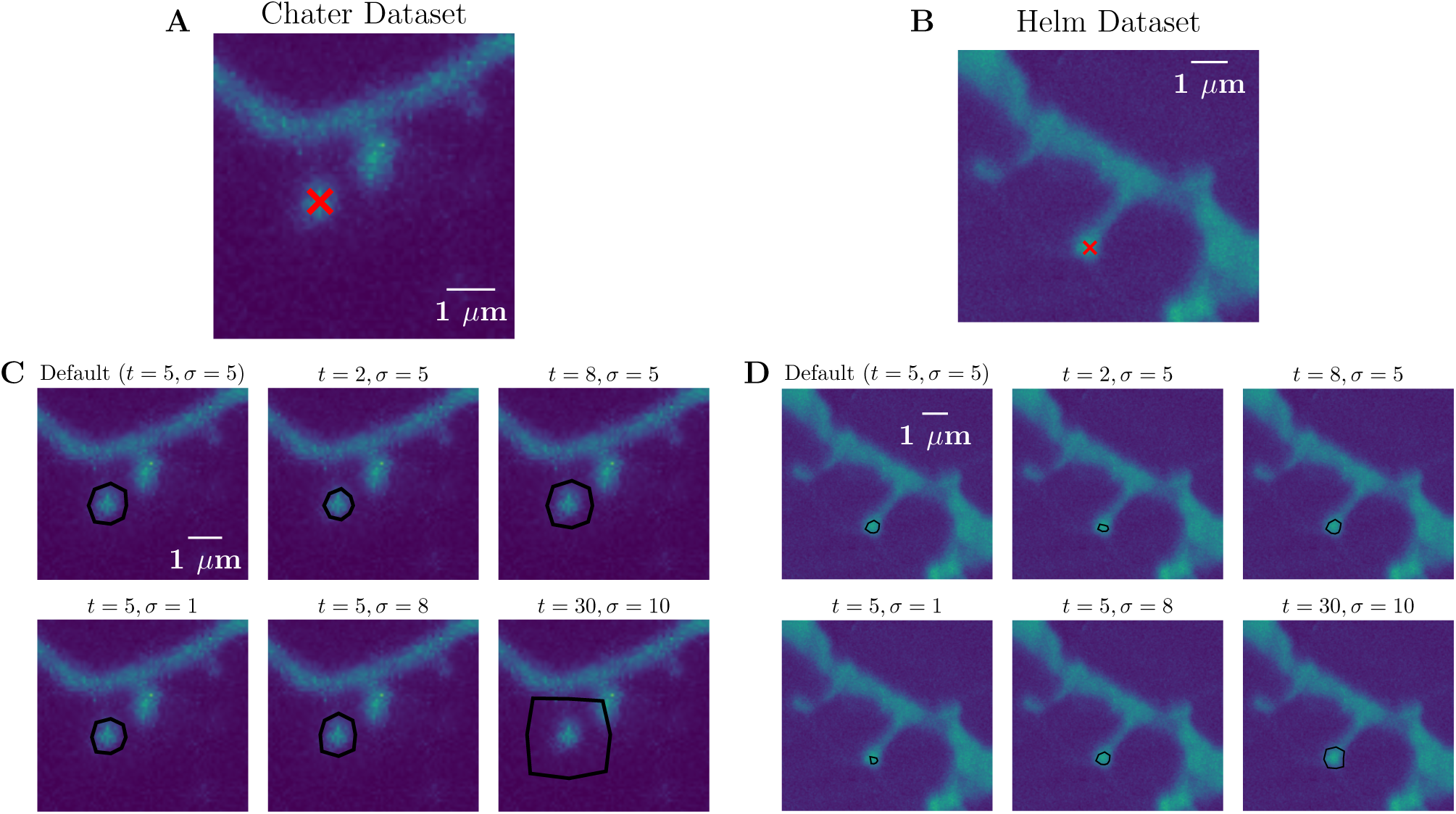
Tuning the parameter settings allows for the automatic ROI generation to adapt to a variety of experimental images. **A)-B)**Example experimental spines marked with a red cross from the Chater et al. (2022) and Helm et al. (2021) datasets, respectively. **C-D)** Given these spines and the marked locations, the code can generate a set of different ROI given different values of the tolerance *t*, and *σ* of the canny edge detection filter. We note that certain parameter combinations work better for certain datasets (e.g., the default setting generates a satisfactory ROI for the Chater spine, while the *t* = 30*, σ* = 10 generates the correct ROI for the Helm spine. All these of ROIs are automatically generated and have not been edited using the editable nodes.

**Figure S6:**
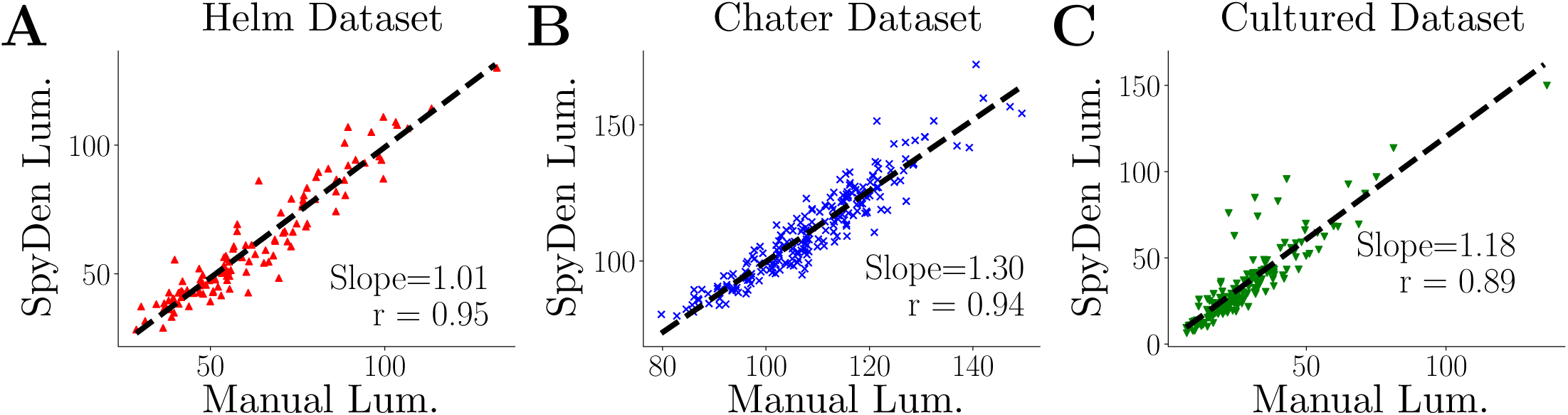
SpyDen agrees across a variety of different experimental conditions. **A-C)** Comparison of the luminosity calculated using the manual ROIs against the luminosity of the SpyDen ROIs for Helm, Chater and Cultured dataset, respectively. In each case, a linear fit is overlaid showing good agreement between the two evaluations.

**Table S5:** output file containing measurements for individual punctum from puncta detection pipeline.

| dend_puncta.csv and soma_puncta.csv |  |
| --- | --- |
| Column Name | Definition |
| Id | Unique identification number assigned to each punctum. |
| channel | Number of channel for which the punctum is located on |
| RoiId | Identification number of the dendritic ROI to which the punctum belongs |
| snapshot | Time point in a time series data to which the punctum belongs |
| location | Pair of x,y coordinate on the image for the punctum |
| radius | Radius of the circular punctum |
| max | mMaximum fluorescent intensity of the punctum |
| min | Minimum fluorescent intensity of the punctum |
| mean | Mean fluorescent intensity of the punctum |
| std | Standard deviation of fluorescent intensities of the punctum. |
| median | Median fluorescent intensity of the punctum |
| distance | Distance from the start of dendritic ROI along the ROI |

**Figure S7:**
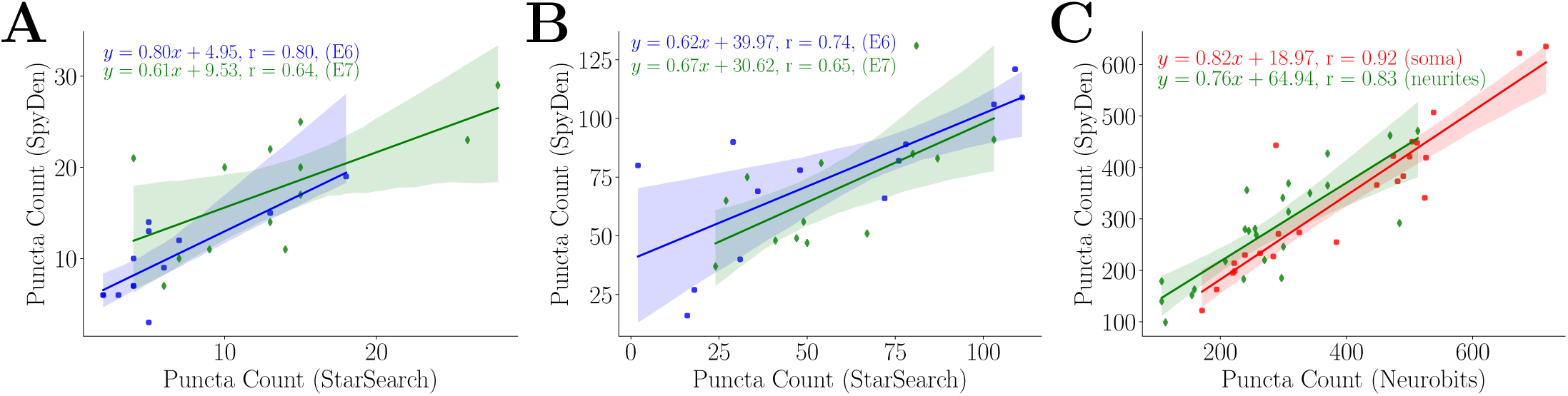
SpyDen puncta detection is in good agreement with other specific-tools across multiple experimental conditions. **A-C)** Comparison of the puncta count in smFISH from (Ciolli Mattioli et al. (2019)) for different isoforms of cdc42 using StarSearch and SpyDen in Neurites (A) and Soma (B) ROIs. C) Comparison of puncta count in FISH images from (Fonkeu et al. (2019)) using SpyDen against Neurobits. In each case, a linear fit is overlaid (with 95% confidence Intervals) showing good agreement between the two evaluations.

### Algorithms

#### Algorithm 1 Finding of the dendritic medial axis path

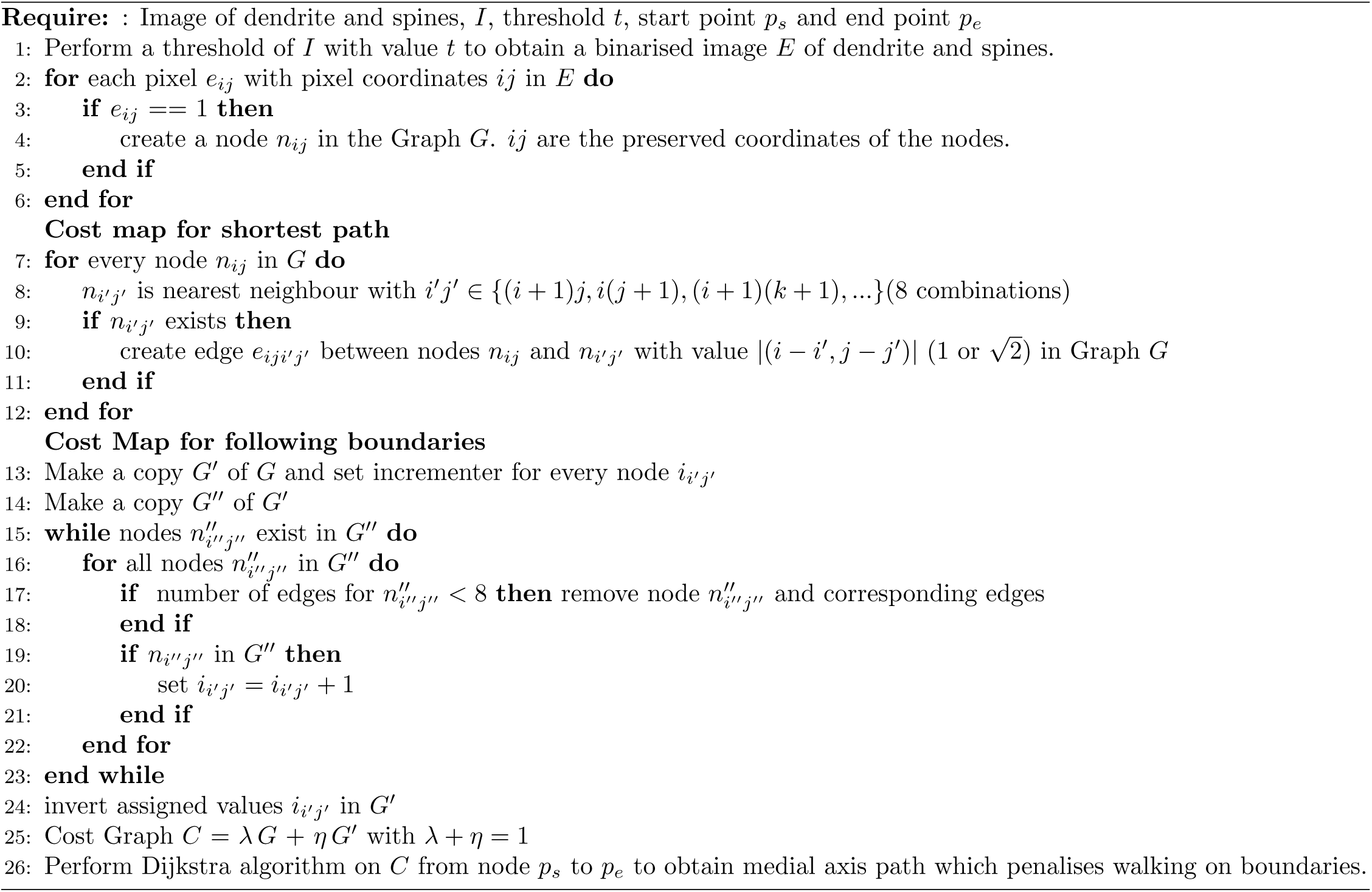

#### Algorithm 2 Finding of the dendritic width

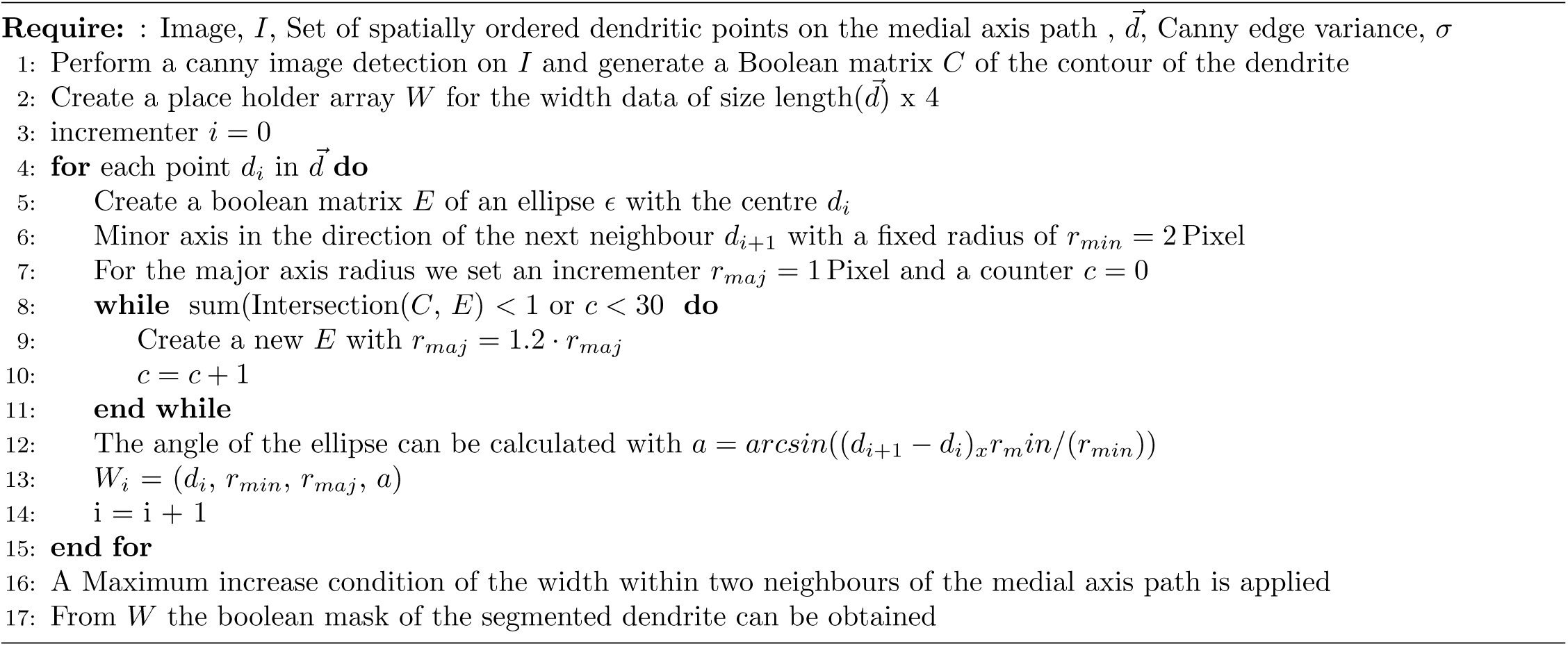

#### Algorithm 3 Finding the region of interest for a spine

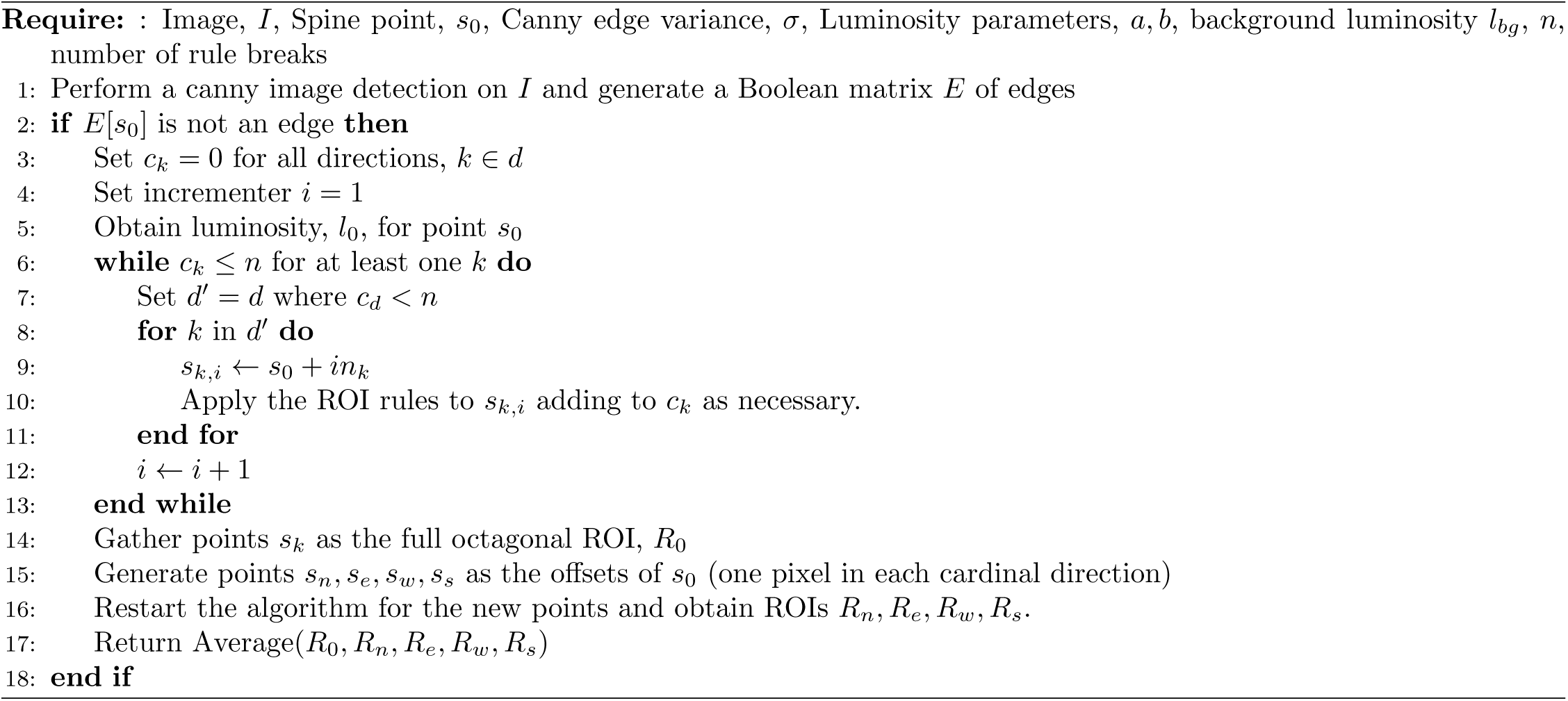

### ROI generation rules

The rules used for the generation of the synapse ROIs (and described briefly in the main text and the algorithm above) are presented here in a detailed manner:

1. **Boundary Rule** If *s_d,i_* exceeds the boundary of the image, the progression in this direction is immediately halted, i.e., *c_d_* is set to be *n*.
2. **Contour Rule** The image is treated with a Canny edge detection algorithm that takes a pre-determined sigma parameter that defines the Gaussian kernel and retrieves edges in the subject matter. If the *s_d,i_* encounters one such edge, this adds a certain number of strikes to *c_d_* depending how far we are from *x_o_*
3. **Luminosity Fall-off Rule** If the luminosity, *l_d,i_* becomes luminosity, *l*_0_*/a* or the luminosity falls below *b* times the background luminosity *l_bg_*, the algorithm adds one to *c_d_*, i.e., you continue while

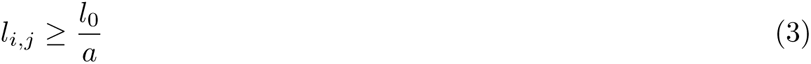

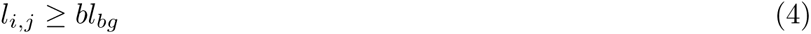
4. **Dendrite Rule** We assume that the spine, on average, is symmetrical and that the user will supply the centre of the spine for (*x*_0_*, y*_0_). Therefore, if *v_i,j_* is closer to the dendrite centre than the initial point, *c_d_* is also incremented.
5. **Symmetry Rule** As a direct consequence of the previous assumption, we also introduce the symmetry rule: the ray directly opposite cannot be more than twice as long before triggering a strike. Assuming that *s_−d,i_* is a ray that has stopped progressing (i.e., *c_−d_ ≥ n*), we can mathematically describe the moment that one is added to *c_d_*

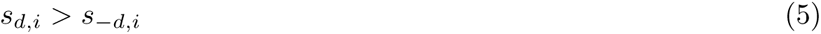
6. **Luminosity Increase Rule** Another assumption of the algorithm is that point *s*_0_ is among the brightest pixels within the spine. Thus, any consistent increase in luminosity is indicative that we have entered the dendrite or another spine. Therefore, persistent luminosity strengthening also leads *c_d_* being increased by one.
7. **Overlap Rule** We note that ROIs of different spines should not overlap, as then some quantities will be counted at least twice. Therefore, given another spine location 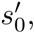 the distance between *s_d,i_* and *s*_0_ must be less than to 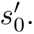

This simple set of rules leads to a small set of parameters for the user to use, meaning that robustness and reproducibility are maintained across a wide array of experimental conditions. These parameters are as follows:

1. The threshold *n* that defines when the ray progression ends
2. The value of variance, *σ* of the Gaussian filter in the canny edge detection
3. The values *a* and *b* of the luminosity thresholds.

The initial values that have worked well for the examples in this article are set to *n* = 3, *σ* = 1.5, *a* = 3 and *b* = 4. However, the option to change these values and turn on and off certain features in the ROI generation allows for a broader use case where certain assumptions or rules may not be valid.

## Notes

https://github.com/meggl23/SpyDen

https://github.com/meggl23/SpydenPaperFigs

https://gin.g-node.org/CompNeuroNetworks/SpyDenTrainedNetwork

### Competing Interest Statement

The authors have declared no competing interest.

